# Genome-Wide Screens Identify Core Regulators of Cell Surface Prion Protein Expression

**DOI:** 10.64898/2025.12.07.692843

**Authors:** Kathryn S. Beauchemin, Surachai Supattapone

## Abstract

Expression of the cellular prion protein, PrP^C^, on the surface of neurons plays an important role in the pathogenesis of prion disease. We performed genome-wide CRISPR/Cas9 knockout screens in prion-infectible cells of neuronal origin (CAD5) to identify regulators of cell surface PrP^C^ expression. We identified and validated 46 positive and 21 negative regulators of cell surface PrP^C^ expression in undifferentiated CAD5 cells. Pathway analysis of the screening dataset showed that genes involved in the glycophosphatidylinositol (GPI) anchor and N-glycosylation biosynthetic pathways were overrepresented as positive regulators of cell surface PrP^C^. We also sought to determine whether the same or different genes regulate cell surface PrP^C^ in CAD5 cells that have been differentiated to a more neuronal state and validated 41 positive and 13 negative regulators of CAD5 cell surface PrP^C^ expression in the differentiated state. We identified 23 core genes as shared between the undifferentiated and differentiated cell states, including many positive regulators involved in GPI anchor biosynthesis. Intriguingly, unique regulators were also identified in the undifferentiated and differentiated cell states, suggesting that some mechanisms regulating cell surface PrP^C^ expression in CAD5 cells are dependent on cell state. This list of core genes involved in regulating cell surface PrP^C^ expression in a prion-susceptible, neuron-like cell type offers a valuable guide for future research and may help identify potential therapeutic targets for prion disease and other neurodegenerative diseases.

## Introduction

Prion diseases are invariably fatal and affect humans and mammals^1–3^. Prions are unconventional pathogens which lack a transmissible nucleic acid genome^4,5^. Infectious prions are formed when the normal, cellularly expressed prion protein, PrP^C^, misfolds into a pathogenic form, PrP^Sc^ ^6^. While PrP^C^ is highly alpha-helical, PrP^Sc^ is rich in beta sheets^7^, is resistant to degradation by proteases^8^, and is prone to aggregation^9^. PrP^Sc^ accumulates in the brain resulting in the neuropathological hallmarks of prion disease including spongiform change, neuronal loss, and gliosis^10^.

PrP^C^ is a 30-40kDa glycoprotein which is attached to the outer leaflet of the plasma membrane by a glycosylphosphatidylinositol (GPI) anchor^11^. PrP^C^ has two sites for N-linked glycosylation and exists in three glycoform states (di-glycosylated, mono-glycosylated, and un-glycosylated) of complex type^12–15^. The amino acid sequence of PrP^C^ is highly conserved^16^ and expression is nearly ubiquitous amongst tissues, with the highest expression levels in nervous tissue^17^. Like other secreted proteins, PrP^C^ is co-translationally translocated on ribosomes at the ER, transits to the Golgi, and is delivered to the cell surface (reviewed in^18^). Most PrP^C^ is localized to lipid rafts at the cell surface^19,20^, though some is transferred to clathrin-coated pits where it undergoes endocytosis and recycling^21,22^. PrP^C^ is also subject to several different endoproteolytic events (reviewed in^23^). The physiological function of PrP^C^ is unclear, but studies have implicated PrP^C^ in various nervous system functions including the maintenance of peripheral nerve myelination, synapse formation, and neuroprotection (reviewed in^24^). Nonetheless, knockout mice lacking PrP^C^ appear generally healthy^25–30^.

Despite significant efforts, to date, there are no clinically approved prophylactics or therapeutics for any type of prion disease (reviewed in^31^). However, even partial reductions in PrP^C^ significantly extend the lifespan of prion-infected animals^32–34^. Recent pre-clinical success using genetic means to reduce PrP^C^ using Zinc Finger Repressors^35,36^, antisense oligonucleotides^33,37,38^, liposomal-siRNA-peptide complexes^39^, divalent siRNA^34^, *in vivo* base editing^40^ and epigenetic modifiers^41^ emphasize the strength of this strategy, though delivery of modifiers to the brain remains a challenge.

Recent studies have also indicated that the oligomeric form of β-amyloid peptide (Aβo), the species responsible for the initiation of Alzheimer’s disease pathology^42,43^, binds cell-surface PrP^C^ with high-affinity^44^. PrP^C^ acts as a neuronal receptor for Aβo and binding is required for toxic Aβo signaling to occur^44–48^, thus preventing this interaction with antibodies^49–53^, small molecules^54^, or with genetic depletion of PrP^C^^44,45,48,55^ rescues Alzheimer’s pathophysiology. Additionally, PrP^C^ interacts with and influences the behavior of other neurodegeneration-associated proteins including tau^56,57^, α-synuclein^58^ and TAR DNA-binding protein 43 (TDP-43)^59^. Therefore, reduction of cell-surface PrP^C^ may be beneficial as a general strategy in the treatment of neurodegenerative disease.

In recent years, genome-wide screens to elucidate the molecular details of the metabolism of PrP^C^ have been performed in a variety of cell types including human haploid leukemic cells^60^, neuroectodermal cells^61^, and glioblastoma cells^36,61,62^. While these efforts have identified several key factors associated with PrP^C^ metabolism, no genome-wide screens for modulators of cell-surface PrP^C^ have been performed in prion-infectable cells. CAD5 cells are a mouse CNS cell line derived from catecholaminergic neuronal cells which can be infected with multiple murine prion strains, can be genetically engineered to be infectable with prion strains from other species, and which are widely used to study prion cell biology^63–80^. Thus, we endeavored to perform genome-wide CRISPR/Cas9 KO screens in CAD5 cells to identify regulators of cell surface PrP^C^ in a prion-infectable cell line.

## Materials and Methods

### Cell lines and cell culture

CAD5 and HEK293FT cell lines were kindly provided by Charles Weissmann (Scripps Florida, Jupiter, FL, USA) and Michael Cole (Geisel School of Medicine at Dartmouth, Lebanon, NH, USA), respectively.

CAD5 cells were maintained in Opti-MEM I Reduced-Serum Media with GlutaMAX (Gibco, Waltham, MA, USA) supplemented with 10% HyClone Bovine Growth Serum (BGS) (Cytiva, Marlborough, MA, USA) and 1X penicillin/streptomycin (Corning, Corning, NY, USA). To differentiate CAD5 cells, cells were plated at desired confluency (60-90%). The next day, cells were gently rinsed 2X with PBS and medium was replaced with DMEM:F12 (Corning) containing 50 ng/mL sodium selenite (Sigma-Aldrich, St. Louis, MO, USA) (protein-free media, PFM). Cells were cultured in PFM for a total of 4 days, with a media refresh at 48 hr.

HEK293FT cells were cultured in IMDM (Gibco) supplemented with 10% HyClone Fetal Bovine Serum (FBS) (Cytiva), 6 mM L-glutamine (Corning), and 1X non-essential amino-acids (MilliporeSigma, Burlington, MA, USA) for the first 24 hr following thawing. After 24 hr, G418 (MilliporeSigma) was added to the growth medium to a final concentration of 500 µg/mL. Cell lines were maintained at 37°C, 5% CO_2_.

Cell lines were routinely monitored for mycoplasma contamination using the LookOut^®^ Mycoplasma PCR Detection Kit (Sigma-Aldrich).

### Plasmids

All plasmids were obtained from Addgene (Cambridge, MA, USA). Mouse Brie CRISPR knockout in lentiGuide-Puro pooled library was a gift from David Root and John Doench (Addgene #73633 ; http://n2t.net/addgene:73633 ; RRID:Addgene_73633). LentiCas9-Blast was a gift from Feng Zhang (Addgene plasmid # 52962 ; http://n2t.net/addgene:52962 ; RRID:Addgene_52962) and lentiGuide-Puro was a gift from Feng Zhang (Addgene plasmid # 52963 ; http://n2t.net/addgene:52963 ; RRID:Addgene_52963). pMD2.G was a gift from Didier Trono (Addgene plasmid # 12259 ; http://n2t.net/addgene:12259 ; RRID:Addgene_12259) and psPAX2 was a gift from Didier Trono (Addgene plasmid # 12260 ; http://n2t.net/addgene:12260

; RRID:Addgene_12260).

### Antibodies and reagents

Blasticidin and puromycin were purchased from Gibco (Thermo Fisher Scientific, Waltham, MA, USA), 1X PBS without calcium or magnesium was obtained from Corning, and polybrene was obtained as a 10 mg/mL stock from MilliporeSigma.

The primary antibodies used in this paper were: anti-PrP 6D11 (mouse monoclonal, 1:20,000) (Covance, catalog# SIG-399810), anti-CD59:APC (recombinant rabbit monoclonal, 2.5uL per 1x10^6^ cells) (Sino Biological, catalog # 12474-R029-A). Rat anti-mouse IgG2a:PE (1uL per 1x10^6^ cells) (Thermo Fisher, catalog # 12-4210-82) was used as a secondary antibody for flow cytometry. Anti-mouse IgG sheep antibody (HRP (Horseradish Peroxidase)) was used as a secondary antibody for Western blots (Cytiva, catalog # 95017-332).

The inhibitors 1-deoxynojirimycin hydrochloride (DNJ), 1-Deoxymannojirimycin hydrochloride (DMJ), and swainsonine (SWA) were purchased from Millipore Sigma and reconstituted at 10 mM in DMSO or _d_H20. HM03 was purchased from MedChemExpress (Monmouth Junction, NJ,

USA) as a 10 mM stock solution in DMSO. All inhibitors were aliquoted, stored at the manufacturer’s recommended temperature, and discarded after use to avoid freeze/thaw.

### Generation of Cas9-expressing CAD5 monoclonal lines

HEK293FT cells were used for lentiviral packaging of LentiCas9-Blast. HEK293FT cells were seeded in 6-well plate (Corning) to achieve 80% confluence on the day of transfection. Media was changed 1 hr before transfection to media containing 2 mM caffeine (Sigma-Aldrich, St. Louis, MO, USA). Transfection was performed using 1ug lentiCas9-Blast, 3uL LipoD293 (Signagen, Rockville, MD, USA), and the plasmids pMD2.G and psPAX. The next day, the media was changed to include 2% FBS and 2 mM caffeine. The following day, the supernatant was collected and 0.45 uM filtered (MilliporeSigma).

The resulting lentiCas9-blast lentiviral stock was used immediately to transduce a 100 mm plate of WT CAD5 cells with 10 µg/mL polybrene. The following day, virus-containing media was replaced with complete media and cells were allowed to recover. Blasticidin selection occurred for the next 6 days, with the addition of media containing 5 µg/mL blasticidin every other day (total of 3 treatments). Individual clones were ringed and grown out as monoclonal lines.

### Amplification of pooled lentiviral library

The Mouse Brie CRISPR knockout pooled library plasmid library was amplified in Endura electrocompetent cells per the manufacturer’s protocol (Lucigen, Middleton, WI, USA). Library was electroporated two times and recovery reactions were rotated at 250 rpm for 1 hr at 37°C. Transformations were pooled and a dilution plate was prepared to calculate transformation efficiency and ensure that each sgRNA construct was represented at least 50X in the amplified library. Aliquots (1 mL) of the pooled reactions were plated on 4 x 500 cm^2^ LB-Ampicillin (100 µg/mL) agar bioassay plates (Thermo Fisher Scientific, Waltham, MA, USA). Plates were incubated at 32 °C for 14 hr. Colonies were harvested by washing each plate with 15 mL LB medium twice using a cell scraper and pooled. Endotoxin Free Plasmid Maxi Kit (Qiagen, Germantown, MD, USA) was used to isolate DNA per the manufacturer’s protocol. Next generation sequencing was used to check for sgRNA representation and evenness.

### Lentivirus production in HEK293FT cells

HEK293FT cells were seeded 24 hr before transfection in 15 cm plates to allow for ∼60% confluence on the day of transfection. Medium containing G418 was removed 30 min before transfection and replaced with 25 mL of IMDM supplemented with 10% FBS, 6 mM L-glutamine, and 1X non-essential amino acids (transfection medium), with 12.5 μg Brie library DNA, 7.5 µg psPAX2, and 5 μg pMD2.G and gently vortexed. LipoD293 (Signagen) (75 μL) was added to 1.25 mL of the transfection medium and gently vortexed. This diluted LipoD293 was immediately added to the DNA mixture, gently vortexed, and incubated for 10 min at room temperature. The LipoD293/DNA mixture was added slowly to the cells with swirling for distribution, then incubated for 24 hr at 37 °C, 5% CO_2_. The medium was then replaced with IMDM supplemented with 2% FBS, 6 mM L-glutamine, 1X non-essential amino acids, and 2 mM caffeine and incubated for an additional 24 hr. Lentiviral particles were harvested by filtration through a 0.45-μm filter and concentrated 10X using Amicon Ultra-15 10kDa Centrifugal Filter units (MilliporeSigma).

### Lentivirus titering and transduction into WT CAD5 cells

The Brie CRISPR knockout pooled library was designed to target 19,674 genes in total. Within the library, each gene is targeted by up to 4 unique sgRNAs. We defined one copy of the library as a 500-fold representation per sgRNA, so a copy of the library is approximately 40 million cells. During culture, passage, and cryopreservation, no fewer than 1 copy with 500-fold representation was always maintained to protect representation.

CAD5 WT cells were seeded 24 hr before transduction in 6-well plates (Corning) to allow for ∼60% confluence on the day of transduction. Serial dilutions (5-fold) of lentiviral particles ranging from 10^-1^ to 5^-3^ were prepared in growth medium containing 8 µg/mL polybrene and 2 mL of the diluted lentiviral particles were added to the cells. After 24 hr, virus and polybrene-containing medium was replaced with growth medium. After 24 hr, selection was initiated by addition of puromycin (3.5 μg/mL final concentration) to the growth medium. Medium was replaced every 48 hr with fresh medium containing 3.5 µg/mL puromycin for a total of 5 days. The lentivirus titer was calculated using the formula:

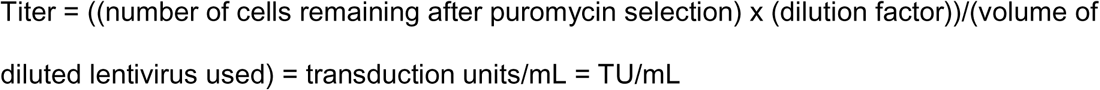

A monoclonal line of CAD5 cells constitutively expressing Cas9 were seeded in 150 mm plates 24 hr before transduction to allow for ∼60% confluence on the day of transduction. Cells were transduced with lentivirus at an MOI of 0.3 in the presence of 8 μg/mL polybrene. The total number of cells that were transduced ensured 500X representation of each single-guide RNA (sgRNA) construct in the Brie library. The medium was replaced with growth medium after 24 hr of incubation at 37°C, 5% CO_2_, and 24 hr later, 3.5 µg/mL puromycin was added for selection of transduced cells. Cells were grown under selection for 5 days with medium renewal every 48 hr. CRISPR/Cas9-modified CAD5 cells were expanded, and to ensure maintenance of adequate representation, library cells were frozen down at 1000X-1500X representation of each sgRNA construct in the library. Library was stored in liquid nitrogen until use. Genomic DNA was extracted from 4 x 10^7^ cells (500X representation) using the Blood and Cell Culture DNA Midi Kit (Qiagen) to check the library for sgRNA representation through next generation sequencing, as detailed below.

### FACS for cell surface PrP^C^

For each replicate in the genome-wide screen, library was thawed at 1000X-1500X representation and cultured for 1 week to allow for recovery from freeze/thaw. For undifferentiated screens, the day before screening, a minimum of 4 x 10^7^ CAD5 CRISPR/Cas9 KO library cells were counted and seeded for growth overnight. For differentiated screens, a minimum of 4 x 10^7^ cells were plated to achieve ∼60% confluence the next day and cultured in protein-free media (PFM) for a total of 4 days, as described above. In both the undifferentiated and differentiated screens, the day of sorting, cells were gently rinsed 1X with PBS, lifted with CellStripper (Corning), and transferred to a tube containing an equal volume of complete growth medium. 4 x 10^7^ cells were aliquoted into one tube and 3 additional aliquots of 1 x 10^6^ cells each were used for staining controls. Cells were spun at 150 x *g* for 5 min at room temperature. The pellet was resuspended in PBS supplemented with 2% FBS (staining buffer). Spinning and resuspension were repeated once more to ensure removal of all media and lifting agent.

4 x 10^7^ library cells were suspended in 5 mL staining buffer and stained with anti-PrP (6D11) antibody at a 1:20,000 dilution. Cells were next stained with rat anti-mouse IgG2a:PE antibody at a concentration of 1uL of stock antibody per 1 x 10^6^ cells. Finally, cells were stained with anti-CD59:APC antibody at a concentration of 2.5 µL stock antibody per 1 x 10^6^ cells. Each sequential staining step included at 45 min incubation at 4 °C in the dark with brief vortexing every 15 min. Following each antibody incubation step, cells were pelleted at 150 x *g* for 5 min at 4 °C, then resuspended in 10 mL staining buffer. The spin and rinse steps were repeated for a total of 3 spins, and the final resuspension volume was 5 mL staining buffer, except for the final resuspension pre-sorting, which was 3 mL. Cells were filtered through a 40 µm cell strainer (Falcon Plastics, Brookings, SD, USA) and stored in the dark, on ice, until sorting.

A FACSAria III Cell Sorter (BD Biosciences, Franklin Lakes, NJ, USA) with a 70 µM nozzle and 488 nm and 635 nm excitation lasers was used for FACS. For each replicate, 4 x 10^7^ viable cells were sorted at a rate of approximately 3000 events/sec. Cells were gated based first on FSC and SSC characteristics to isolate single cells. Next, cells were gated based on CD59 (APC) fluorescence (middle 90% fluorescence), then subgated based on PrP (PE) fluorescence. Cells within the lowest 20% and highest 20% PrP fluorescence were collected and called “PrP Low” and “PrP High”, respectively. Selection with this gating strategy yielded approximately 4 x 10^6^ output cells in each subset (PrP Low and PrP High) per screening replicate, which were immediately processed for genomic DNA extraction.

### Sequencing and analysis of sgRNA enrichment

Genomic DNA was isolated from cells collected via FACS using the Blood and Cell Culture DNA Midi Kit (Qiagen) per manufacturer’s instructions. Three subsequent rounds of PCR were performed to amplify the gDNA region containing the sgRNA lentiviral cassette, to attach Illumina sequencing adaptors and barcodes, and to clean up PCR products. Primers are listed in Supplementary Tables, “PCR Primers” tab.

All PCR reactions were performed using Q5 Hot Start High Fidelity 2X Master Mix (NEB, Ipswich, MA, USA) as per manufacturer’s recommendations with a reaction volume of 50 µL and 25 amplification cycles.

PCR1 amplified sgRNA sequences from genomic DNA using common flanking sequences. PCR1 reactions were performed to consume all genomic DNA, using 1.5 µg gDNA per reaction. PCR1 products were pooled and purified using a QIAquick PCR Purification Kit (Qiagen).

PCR2 attached Illumina adaptors and barcodes. Uniquely barcoded forward primers were used for each sample for facile sample identification and a pooled mix of twelve PCR2 reverse primers were used for each sample to increase diversity. 8 reactions were performed per sample containing 10 µL of pooled and purified PCR1 product as input.

PCR3 cleaned up amplicon ends to improve clustering on the sequencer. 8 reactions were performed per sample using 150 ng of pooled and purified PCR2 product as input. PCR3 product was pooled and run on a 1.5% OmniPur agarose (MilliporeSigma) gel prior to gel extraction using a QIAquick Gel Extraction Kit (Qiagen). Gel extracted product was concentrated using a QIAquick PCR Purification Kit (Qiagen).

### Next Generation Sequencing

All sequencing was performed by the Dartmouth Genomics Shared Resource core facility. Fragment Analyzer (Agilent Technologies, Santa Clara, CA, USA) and DNA quantification via Qubit Flex (Invitrogen, Waltham, MA, USA) were run as quality control measures prior to NGS runs. An Illumina NextSeq500 (Illumina Inc., San Diego, CA, USA), high output 75 run with single end reads was used for the undifferentiated genome-wide samples and library control, an Illumina NextSeq 2000 P2 1 x 100 run was used for the differentiated genome-wide screening samples and library control, while an Illumina MiniSeq high output 1 x 150 cycle run was used for the secondary library samples. All runs contained a 10% Phi-X (Illumina Inc.) spike in to increase diversity. Read count was minimally 50 reads per sgRNA. Samples were deconvoluted to FASTQ files for subsequent analysis.

### Data Processing and Analysis

Deconvoluted FASTQ files were analyzed using the MAGeCK-VISPR^81^ and MAGeCKFlute pipelines^82^. For the primary genome wide screens, samples were analyzed as biological replicates, comparing enrichment of sgRNAs between the PrP Low and PrP High subsets for each replicate.

To formulate a list of the strongest candidate genes for validation, data from the 3 genome-wide replicate screens (undifferentiated and differentiated) were analyzed as described above. To maximize the scope of follow-up validation and because we were not technically limited by the number of genes to include in a custom secondary validation library, genes with an FDR of ≤ 0.5 were prioritized for validation. Additionally, KEGG analysis was performed and a hand-curated list of genes involved in pathways exhibiting enrichment were selected for inclusion in the secondary library^83–85^.

The secondary library contained 8 unique sgRNAs targeting each gene (Supplementary Tables, “Targeting Secondary Library” tab). To account for the discrepancy between the number of sgRNAs for targeting genes and individual sgRNAs for non-targeting controls for MAGeCK analysis, non-targeting control sgRNAs were randomly assigned gene names, with 10 non-targeting guides per non-targeting gene name (Supplementary Tables, “Non-Targeting Secondary Library” tab). The secondary screens in undifferentiated cells had 3 replicate sorts and the screens in differentiated cells had 5 replicate sorts. Both the undifferentiated and differentiated screens were analyzed using MAGeCK-VISPR and MAGeCKFlute as paired replicates to compare the PrP Low and PrP High samples directly.

Raw .fastq files from all screening samples have been deposited with Mendeley Data and can be found using the following DOIs: 10.17632/jv32cvpn3r.1, 10.17632/5pfdty7zd6.1, 10.17632/p5t6d78zn8.1, 10.17632/vs9fwhbcvj.1, 10.17632/tmkn3zsjts.1, 10.17632/jb89dp34d8.1

### Secondary Library Creation

Non-targeting and targeting secondary libraries were synthesized, amplified, packaged, and transduced into target cells separately. The resulting non-targeting and targeting secondary libraries were mixed 1:1 for use in secondary validation screening.

Custom sgRNA libraries were synthesized as oligo pools by Twist Bioscience (South San Francisco, CA, USA). The oligo pools were amplified as two PCR reactions per library. Each reaction used Q5 Hot Start High Fidelity 2x Master Mix, approximately 13 ng of DNA, 12.5 ng forward, 12.5 ng reverse primers (Supplementary Tables, “PCR Primers” tab). 10 amplification cycles were used to prevent overamplification, per manufacturer’s recommendations. PCR product was pooled and purified using QIAquick PCR Purification Kits.

Golden Gate Assembly was performed using the Golden Gate Bsmb1 Assembly Kit (NEB) to ligate amplified library oligos into the lentiGuide-Puro vector (Addgene # 52963). Resulting plasmids were transformed into One Shot MAX Efficiency DH5alpha-T1 chemically competent cells (Thermo Fisher) and incubated overnight at 37 °C. Colonies were collected, pooled, and plasmids were purified using an EndoFree Plasmid Maxi Kit (Qiagen). Aliquoted plasmid was stored at -80 °C.

Secondary library plasmid was packaged into lentivirus using the same HEK239FT packaging system and titered into CAD5 cells as described for the Brie whole-genome library. The same monoclonal line of CAD5 cells constitutively expressing Cas9 was transduced at an MOI of 0.3, as above, and NGS was used to ensure adequate representation and evenness of sgRNAs.

### Secondary library FACS for cell surface PrP^C^

For each replicate in the secondary validation, library was thawed at 1000X representation and cultured for 1 week to allow for recovery from freeze/thaw. Differentiated and undifferentiated cells were handled as described for the genome-wide FACS screens. On the day of sorting, cells were gently rinsed 1X with PBS, lifted with CellStripper, and transferred to a tube containing and equal volume of complete media for counting. 1.5 x 10^7^ cells were used for each replicate sort, as well as 1 x 10^6^ cells for staining controls. Cells were spun at 150 x *g* for 5 min at room temperature. The pellets were resuspended in PBS supplemented with 2% FBS (staining buffer). Spinning and resuspension were repeated once more to ensure removal of all media and lifting agent.

1.5 x 10^7^ secondary library cells were suspended in 1.875 mL staining buffer and stained with anti-PrP (6D11) antibody at a 1:20,000 dilution. Next, cells were stained with rat anti-mouse IgG2a:PE antibody at a concentration of 1 µL stock antibody per 1 x 10^6^ cells. Each sequential staining step included a 45 min incubation at 4 °C in the dark with brief vortexing every 15 min. Following each antibody incubation step, cells were pelleted at 150 x *g* for 5 min at 4 °C, then resuspended in 10 mL staining buffer. The spin and rinse steps were repeated for a total of 3 spins, and the final resuspension volume was 5 mL staining buffer, except for the final resuspension pre-sorting, which was 3 mL. Cells were filtered through a 40 µm cell strainer (Falcon Plastics, Brookings, SD, USA) and stored in the dark, on ice, until sorting.

An SH800 cell sorter (Sony Biotechnology, San Jose, CA, USA) with a 488 nm excitation laser was used for each replicate sort of the secondary library. For each replicate, 1 x 10^7^ viable cells were sorted at a rate of approximately 2000 events/sec through a 100 µm sorting chip. Cells were gated first based on FSC and SSC characteristics to isolate single cells. Next, cells were gated based on PrP (PE) fluorescence. Cells within the lowest 20% and highest 20% PrP fluorescence were collected and called “PrP Low” and “PrP High”, respectively. Selection with this gating strategy yielded approximately 2 x 10^6^ output cells in each subset (PrP Low and PrP High) per screening replicate, which were immediately processed for genomic DNA extraction.

### Flow cytometry for surface PrP^C^

All flow cytometry experiments were performed on a CytoFLEX flow cytometer (Beckman Coulter, Brea, CA, USA).

Each sample contained approximately 500,000 cells in 100 μL PBS/2% FBS. Primary incubations were performed at 4 °C for 1 hr, followed by three washes in PBS, with final resuspension to 100 µL in PBS/2% FBS. Secondary incubations were performed at 4 °C for 30 min, followed by three washes in PBS, with final resuspension in 250 μL PBS/2% FBS. FlowJo software (BD Biosciences) was used for all flow cytometry analysis. Sequential gating was performed to identify single cell populations. Median fluorescence intensities were used for calculation of percent reduction in surface PrP^C^ expression.

### N-glycosylation inhibitor treatments

Unless otherwise noted, all experiments involving cell treatment with N-glycosylation inhibitors underwent treatment at the indicated final concentration in complete cell culture media, with media/drug refreshes every 48-72 hr.

### Surface protein biotinylation

WT CAD5 cells were grown in 6-well plates and treated with N-glycosylation inhibitors or vehicle controls as described above for 72 hr with media/drug refresh at 48 hr. Cells were washed 3X with 1 mL ice cold PBS. EZ-Link Sulfo-NHS-SS-Biotin (Thermo Fisher) was diluted to 0.5 mg/mL in PBS. 600 µL of EZ-Link Sulfo-NHS-SS-Biotin solution was added to each well and plates were incubated for 1 hr at 4 °C. Following incubation, the biotin solution was aspirated and cells were washed 1X with 1 mL of 25 mM Tris, pH 8.0. Cells were then centrifuged at 150 *x g* for 5 min at 4 °C, resuspended in 1 mL PBS, and washed two additional times in PBS. Cell pellet was finally resuspended in 250 µL lysis buffer (50 mM Tris, pH 8.0, 150 mM NaCl, 0.5% (w/v) sodium deoxycholate, and 0.5% (v/v) Nonidet P-40).

### Phosphoinositide phospholipase (PIPLC) treatment

WT CAD5 cells were grown in 6-well plates and treated with N-glycosylation inhibitors or vehicle controls as described above for 72 hr with media/drug refresh at 48 hr. Cells were washed 3X with 1 mL ice cold PBS, lifted using 500 µL per well enzyme-free dissociation agent, and transferred to 1.5 mL centrifuge tubes. Cells were centrifuged at 150 *x g* for 5 min at 4 °C to pellet cells. Cell pellets were resuspended in 200 µL cold PBS containing 0.25 U/mL PIPLC (Thermo Fisher). Cells were incubated at 37 °C for 1 hr with gentle shaking. Cells were then centrifuged at 150 *x g* for 5 min at 4 °C and resuspended in 1 mL PBS. Spin step was repeated for total of two washes in 1mL PBS each time. Cell pellet was finally resuspended in 250 µL lysis buffer (50 mM Tris, pH 8.0, 150 mM NaCl, 0.5% (w/v) sodium deoxycholate, and 0.5% (v/v) Nonidet P-40).

### Cell lysis and immunoblotting

To immunoblot for PrP^C^, lysed cell solution was centrifuged at 250 x *g* for 30 s to pellet DNA. Lysate supernatant was transferred to fresh 1.5 mL centrifuge tubes. Protein concentrations were determined using a Pierce BCA Protein Assay Kit (Thermo Fisher Scientific) and 10-30 µg total protein was loaded per lane.

SDS-polyacrylamide gel electrophoresis (PAGE) was performed using 1.5 mm 12% polyacrylamide gels with an acrylamide/bisacrylamide ratio of 29:1. The gel was transferred to a methanol-charged polyvinylidene difluoride membrane (MilliporeSigma, Burlington, MA, USA) using a Transblot SD semidry transfer cell (Bio-Rad Laboratories, Hercules, CA, USA). The transfer was set at 2.5 mA/cm^2^ for 45 min. To visualize PrP signal, the membrane was blocked in 5% (w/v) nonfat dry milk (Nestlé, Vevey, Switzerland) in TBST (10 mM Tris, pH 7.1, 150 mM NaCl, 0.1% Tween 20) for 1 h at 4°C. The blocked membrane was then incubated overnight at 4°C with primary antibody, washed three times for 10 min in TBST, then incubated for 1 h with horseradish peroxidase-labeled secondary antibody. Membrane was then washed four additional times for 10 min each in TBST. Blots were developed with SuperSignal West Femto (Thermo Fisher Scientific) chemiluminescence substrate, and images were captured digitally using an Azure 600 (Azure Biosystems, Dublin, CA, USA) imaging system. Relative molecular masses were determined by comparison to PageRuler Plus Prestained Protein Ladder (Thermo Fisher Scientific).

### Cell viability assay

WT CAD5 cells were treated with N-glycosylation inhibitor or vehicle control for 72 hr. Following treatment, cells were rinsed 1X with PBS, lifted in PBS, counted using a Countess 3 automated cell counter (Thermo Fisher), and 1 million cells per sample were transferred to fresh tubes.

Live-or-Dye 405/452 fixable membrane impermeant cell viability dye (Biotium, Fremont, CA, USA) was diluted, aliquoted, and stored per manufacturer’s instructions. 1 µL dye was added to each sample tube, vortexed, and incubated at 4°C for 30 min in the dark. Following incubation, cells were washed 2X with PBS, filtered through a 40 µm cell strainer (Falcon Plastics) and stored in the dark on ice until flow cytometric analysis. To prepare positive control (dead) cells, 1 million cells were incubated at 56 °C for 45 min, allowed to cool to room temperature, and stained using the above protocol. Gating was performed to isolate single cells as described above and positive control (dead) cells were used to identify percentage dead cells in samples. Biological replicate measurements were obtained (n=3) and statistical significance was determined using unpaired t-tests using GraphPad Prism version 10.2.1 for Windows (GraphPad Software, Boston, MA, USA).

### Hspa5 inhibitor treatments

Unless otherwise noted, all experiments involving cell treatment with the Hspa5 inhibitor HM03 were performed by incubating cultured cells in complete cell culture media or protein-free cell differentiation media (PFM) with a final concentration of 5 µM HM03 for 24 hr.

## Acknowledgements

The authors wish to thank Elisabeth Sergison for creating the Cas9-expressing CAD5 monoclonal line, Gary Ward for FACS of the genome-wide libraries, Chris Shoemaker for use of the Sony SH800 sorter as well as technical and scientific guidance, Margaret Ackerman for scientific guidance, Tamutenda Chidawanyika and Kenneth Mark for guidance with CRISPR/Cas9 screening procedures and bioinformatic analysis, Lisa Francomacaro, Rachel Pepin, Abigail Schwind, and Francesca Salerno for scientific guidance and support. Kathryn Beauchemin would also like to thank her husband, Marc Beauchemin, for his unwavering support.

## Conflicts of Interest

K.S.B. and S.S. are co-inventors on a patent application related to the work in this manuscript.

## Funding

This study was funded by the National Institute for Neurological Diseases and Stroke (1R37NS125431, R01NS117276 and R01NS118796 to S.S.) and the National Institutes of Health (5T32AI007519-22 to Deborah Hogan) (P20-GM113132 to Dean Madden).

## Results

### Genome-wide knockout screens reveal genes involved in the regulation of PrP^C^ surface expression in undifferentiated CAD5 cells

To identify genes responsible for the regulation of cell-surface PrP^C^ in a prion-susceptible cell type, we performed genome-wide pooled CRISPR/Cas9 knockout (KO) library screens in the mouse neuronal-like cell line, CAD5. The CRISPR/Cas9-modified CAD5 KO library was generated by first transducing Cas9 into WT CAD5 cells and isolating monoclonal lines (CAD5-Cas9). Next, the Brie mouse whole-genome pooled KO sgRNA library was transduced into CAD5-Cas9 monoclonal cells at a low multiplicity of infection to generate the genome-wide CAD5 KO library^86^**(Figure 1A)**.

**Fig. 1.**
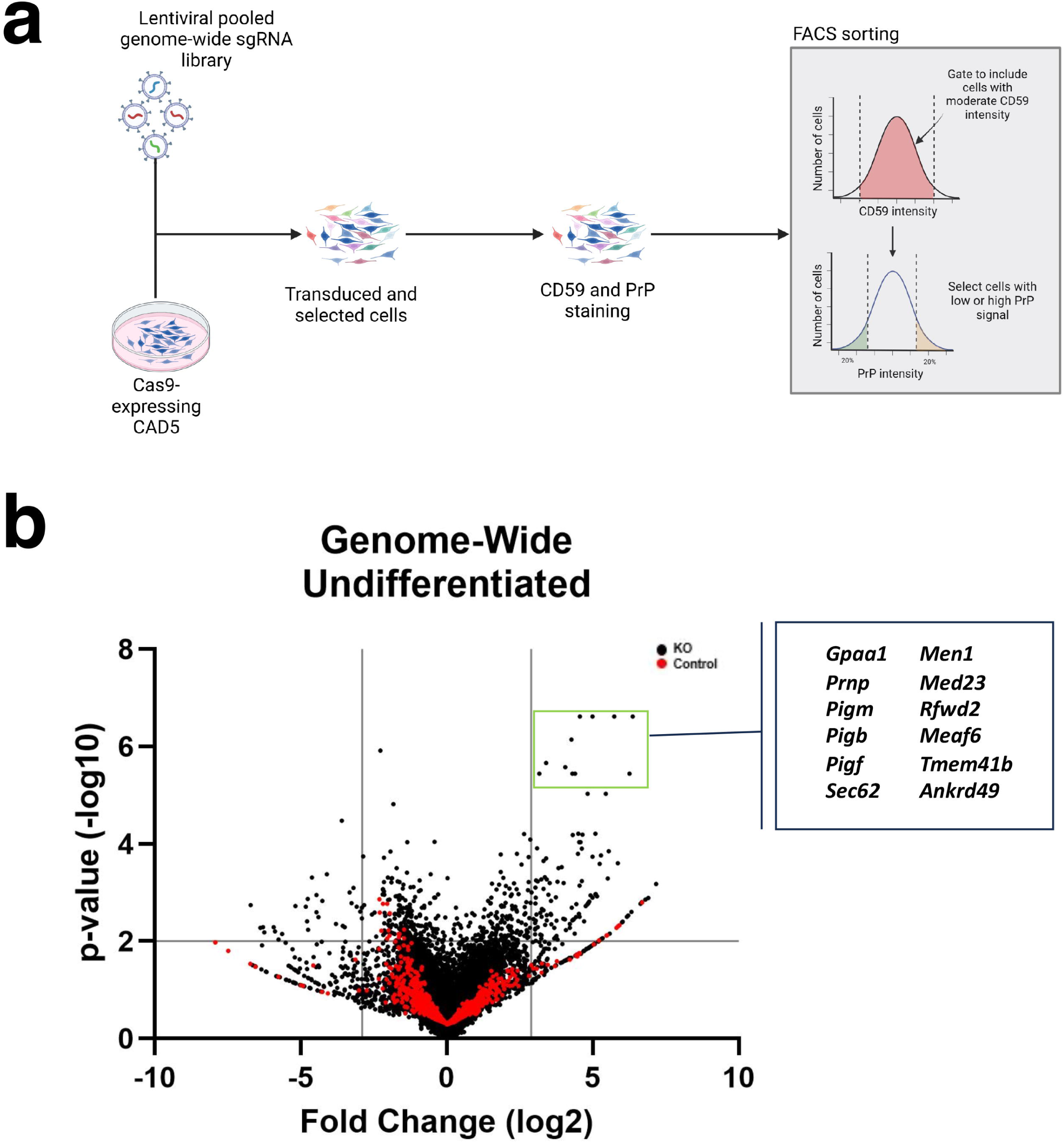
Genome-wide CRISPR-Cas9 screen for positive and negative regulators of PrP^C^ expression at the cell surface of CAD5 cells. (A) Schematic of the CRISPR-Cas9 FACS-based genome-wide screen. (B) Volcano plot of genome-wide KO screen in undifferentiated CAD5 cells with top positive regulators of surface PrP^C^ labeled. Cutoff values indicated with gray lines. Cutoffs were -log_10_(*p*-value) ≥ 2 (corresponding to *p*-value ≤ 0.01) and log fold change ≤ -2.9 or ≥ 2.9 (corresponding to log fold changes outside 3 standard deviations of the mean). KO genes are plotted as black circles and control genes are plotted as red circles.

We performed whole-genome KO screens in biological triplicate. For each replicate screen, the genome-wide CAD5 KO library was doubly stained with antibodies directed at the GPI-anchored cell surface proteins CD59 and PrP and cells were sorted by fluorescence activated cell sorting (FACS). Relative cell surface levels of CD59 were measured via the median fluorescence intensity (MFI) within the allophycocyanin (APC) channel, while relative cell surface levels of PrP^C^ were measured via the MFI within the phycoerythrin (PE) channel. To identify genes specific to PrP^C^ cell surface expression, rather than the expression of all GPI-anchored proteins, CAD5 KO library cells were first gated to exclude cells with exceptionally low or high CD59 expression. Cells within the middle 90% of the CD59 fluorescence histogram were sorted the basis of cell surface PrP^C^. Cells with the lowest or highest 20% of the PrP^C^ fluorescence histogram were sorted into separate populations (PrP^C^ low and PrP^C^ high, respectively) and retained for DNA isolation and next-generation sequencing (NGS) **(Figure 1A, Supplemental Figure 1A)**.

The MAGeCK-VISPR bioinformatic pipeline was used to identify gene hits from the NGS dataset. MAGeCK-VISPR compared the relative sgRNA abundance between the PrP^C^ high and PrP^C^ low populations, resulting in a ranked list of genes based on a robust ranking aggregation (RRA) score with associated significance scores (*p* values) and false discovery rate (FDR) values. The RRA score assigned by MAGeCK-VISPR encompasses the strength of enrichment of individual sgRNAs, degree of consistency seen in enrichment of multiple sgRNAs targeting a specific gene, and agreement across replicate screens **(Supplemental Figure 1B, Supplemental Figure 1C).**

119 genes were identified as positive regulators of CAD5 PrP^C^ surface expression and 29 genes were identified as negative regulators of PrP^C^ surface expression **(Figure 1B, Supplementary Tables, “Genome Undif” tab)**. Inclusion criteria for hits required a log fold change that fell outside 3 standard deviations of the mean (≤ -2.9 or ≥ 2.9) and a *p* value less than or equal to 0.01. *Prnp*, the gene encoding PrP^C^, was a high-ranking positive regulator (hit #2) **(Supplemental Figure 1B)**, lending confidence to our screening dataset.

To validate the hits identified in the whole-genome KO screens, we created a custom secondary KO library comprised of sgRNAs targeting the highest-ranking positive and negative regulators of PrP^C^ surface expression from the whole-genome screens as identified by MAGeCK-VISPR analysis of biological replicates (∼1100 genes total). **(Figure 2A)**. We also included sgRNAs targeting a hand-curated list of genes including gene hits from published PrP^C^ screens and genes which fell within identified enriched pathways from MAGeCKFlute analysis, as well as 2000 non-targeting control guides. To increase the power of the secondary screens, each gene was targeted by eight unique sgRNAs. Targeting sgRNAs for each gene included the same four sgRNAs found in the Brie genome-wide library^86^, two from another genome-wide targeting library called Gouda^87^, and two randomly selected from the GeCKO mouse v2 library^88^. The custom secondary sgRNA library was transduced into CAD5-Cas9 cells and the resulting secondary KO library was sorted in biological triplicate based on PrP^C^ cell surface expression using FACS **(Figure 2A)**.

**Fig. 2.**
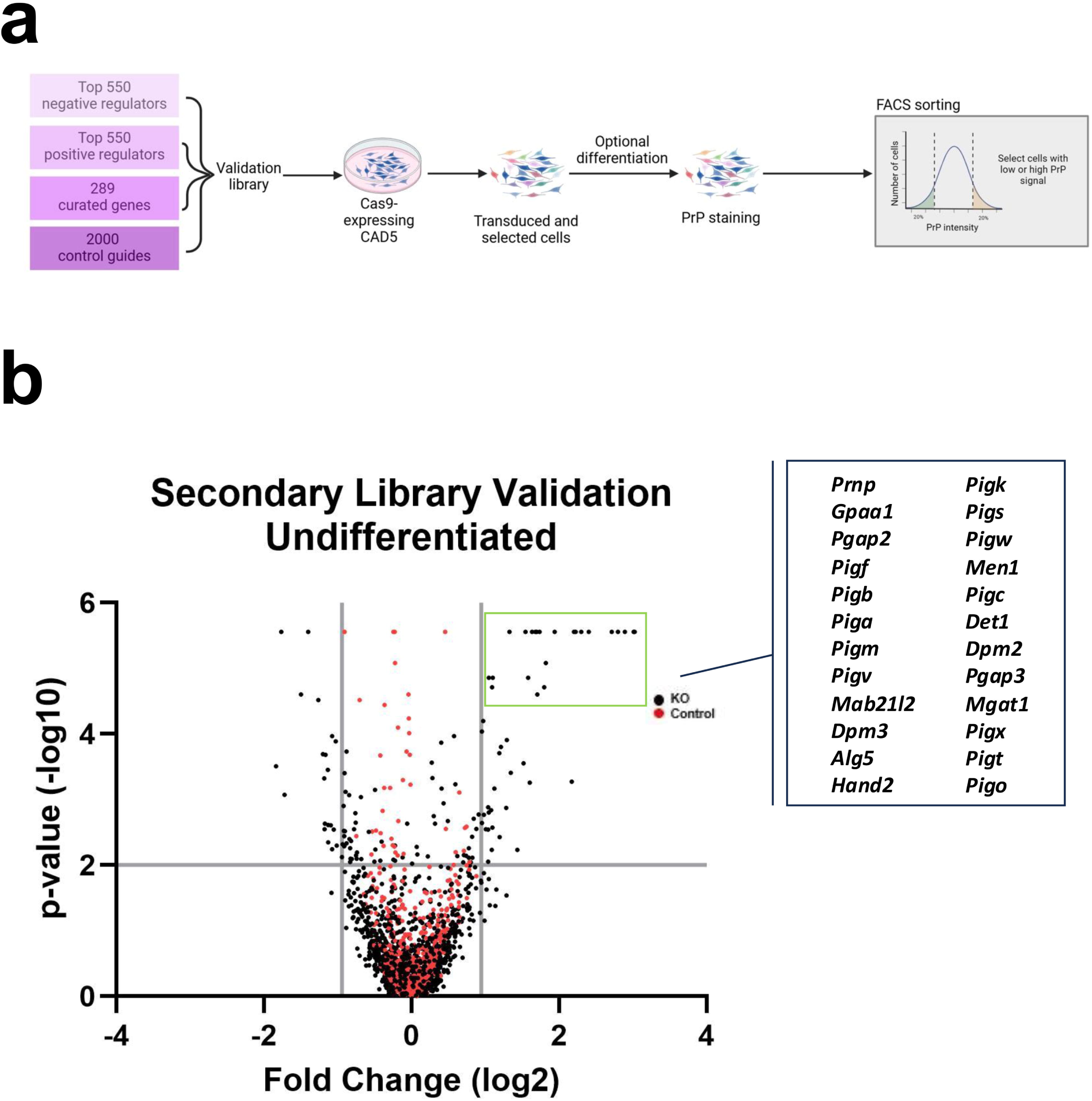
Validation of genome-wide screen for positive and negative regulators of cell surface PrP^C^ expression. (A) Schematic of the CRISPR-Cas9 FACS-based secondary library validation screen. (B) Volcano plot of secondary library validation screen in undifferentiated CAD5 cells with top positive regulators of surface PrP^C^ labeled. Cutoff values indicated with gray lines. Cutoffs were -log_10_(*p*-value) ≥ 2 (corresponding to p-value ≤ 0.01) and log fold change ≤ -0.93 or ≥ 0.93 (corresponding to log fold changes outside 2 standard deviations of the mean). KO genes are plotted as black circles and control genes are plotted as red circles.

Analysis of the resulting NGS data was performed using MAGeCK-VISPR. Hit inclusion criteria for validation of target genes in the secondary library required a log fold change that fell two standard deviations outside of the mean (≤ -0.93 or ≥ 0.93) and a *p* value of less than or equal 0.01. We validated 46 positive regulators and 21 negative regulators of PrP^C^ surface expression **(Figure 2B, Supplementary Tables, “Secondary Val Undif” tab)**. Our secondary validation screens identified *Prnp* as the #1 positive regulator of PrP^C^ surface expression. Taken together, our genome-wide CRISPR/Cas9 KO screens provide a comprehensive list of genes involved in regulation of PrP^C^ at the cell surface of CAD5 cells.

### Bioinformatic analyses identify biosynthetic pathways involved in the regulation of PrPC surface expression

To further dissect the regulation of PrP^C^ utilizing our whole-genome and validation library datasets, we utilized the pathway analysis function of MAGeCKFlute using KEGG, GOBP, REACTOME, and Complex databases **(Figure 3A).**

**Fig. 3.**
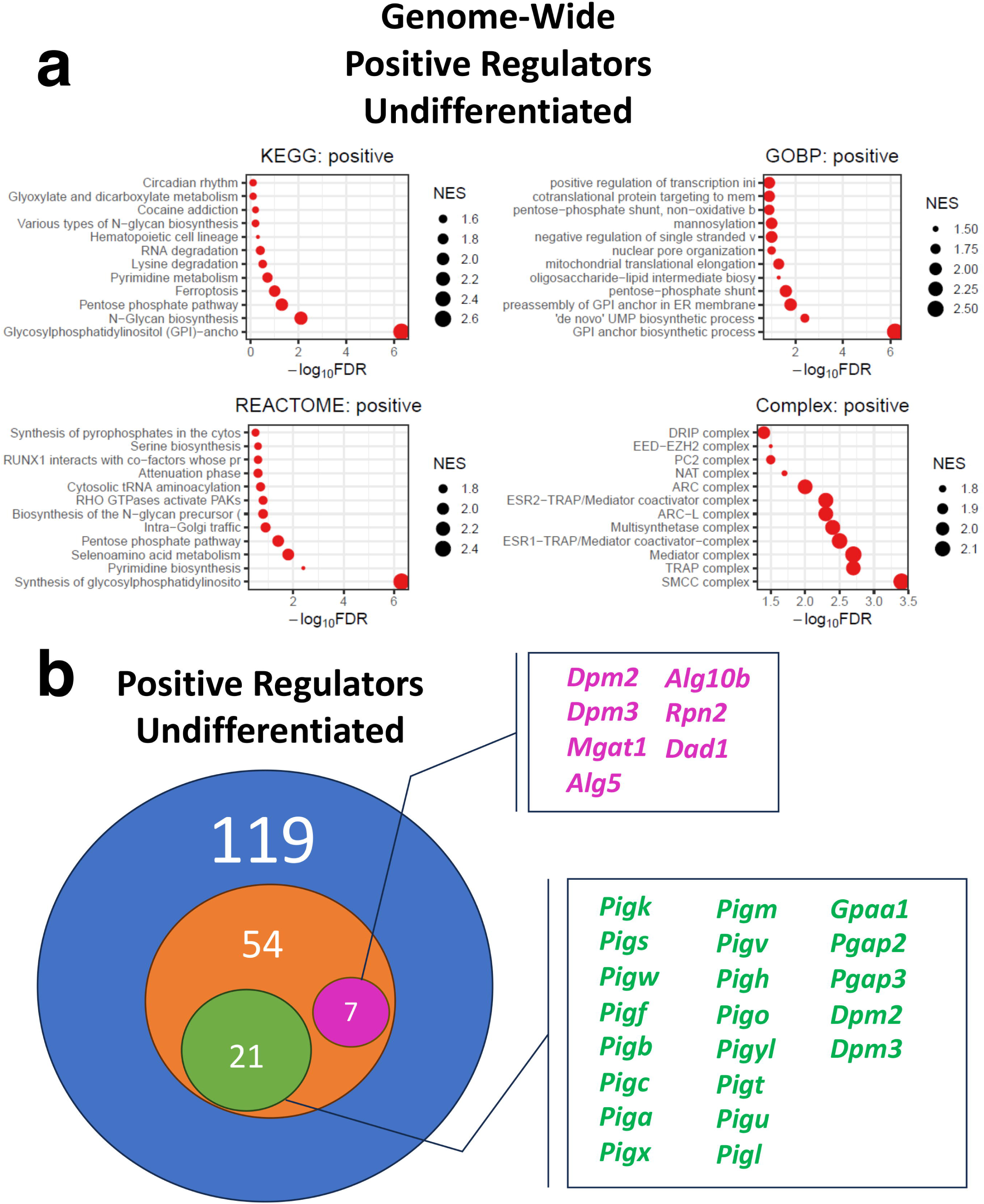
MAGeCKFlute analysis of NGS results from the PrP^C^ cell surface whole-genome knockout screen for pathways which regulate PrP^C^ surface expression in undifferentiated CAD5 cells. (A) Pathway analysis of genome-wide undifferentiated positive hits was conducted using the Kyoto Encyclopedia of Genes and Genomes (KEGG)^83–85^, Gene Ontology Biological Process (GOBP), Reactome, and Complex databases through the MAGeCKFlute platform. Negative log_10_ false discovery rates (FDR) were plotted for pathway hits and point size corresponds to relative normalized enrichment score (NES). (B) Visual diagram comparing the number of positive regulator gene hits identified in the undifferentiated state. The blue circle represents the number of hits identified in the genome-wide screen and the orange circle represents the number of hits identified in the secondary validation screens. Green and purple circles indicate the number of validated genes belonging to the GPI-anchor biosynthesis and N-glycan biosynthesis pathways, respectively. Validated genes belonging to the N-glycan biosynthesis pathway are labeled in purple and GPI-anchor biosynthesis pathway genes are labeled in green.

Unexpectedly, the N-glycan biosynthetic pathway was identified as a positive regulator of PrP^C^ surface expression in the genome-wide screen **(Figure 3A)** and 7 N-glycosylation biosynthetic pathway genes were validated via secondary library screening **(Figure 3B, purple).** We also identified and validated the GPI-anchor biosynthetic pathway as an impressive regulator of PrP^C^ surface expression, as previously reported by Davis et al.^60^**(Figure 3A**, **Figure 3B, green)**. These results pinpoint and underscore the importance of key pathways in PrP^C^ cell surface regulation.

### N-linked glycosylation biosynthetic pathway inhibitors reduce PrPC surface expression in undifferentiated CAD5 cells

To further validate the previously unreported role of the N-glycosylation biosynthetic pathway as a positive regulator of PrP^C^ surface expression, we utilized an orthogonal approach. The N-linked glycosylation biosynthetic pathway includes several residue modification steps which are carried out by distinct enzymes and can be inhibited by small molecules **(Figure 4A)**. We treated CAD5 cells by adding small molecule inhibitors of N-glycosylation enzymes to the culture media and measured the impact on PrP^C^ surface expression via flow cytometry.

**Fig. 4.**
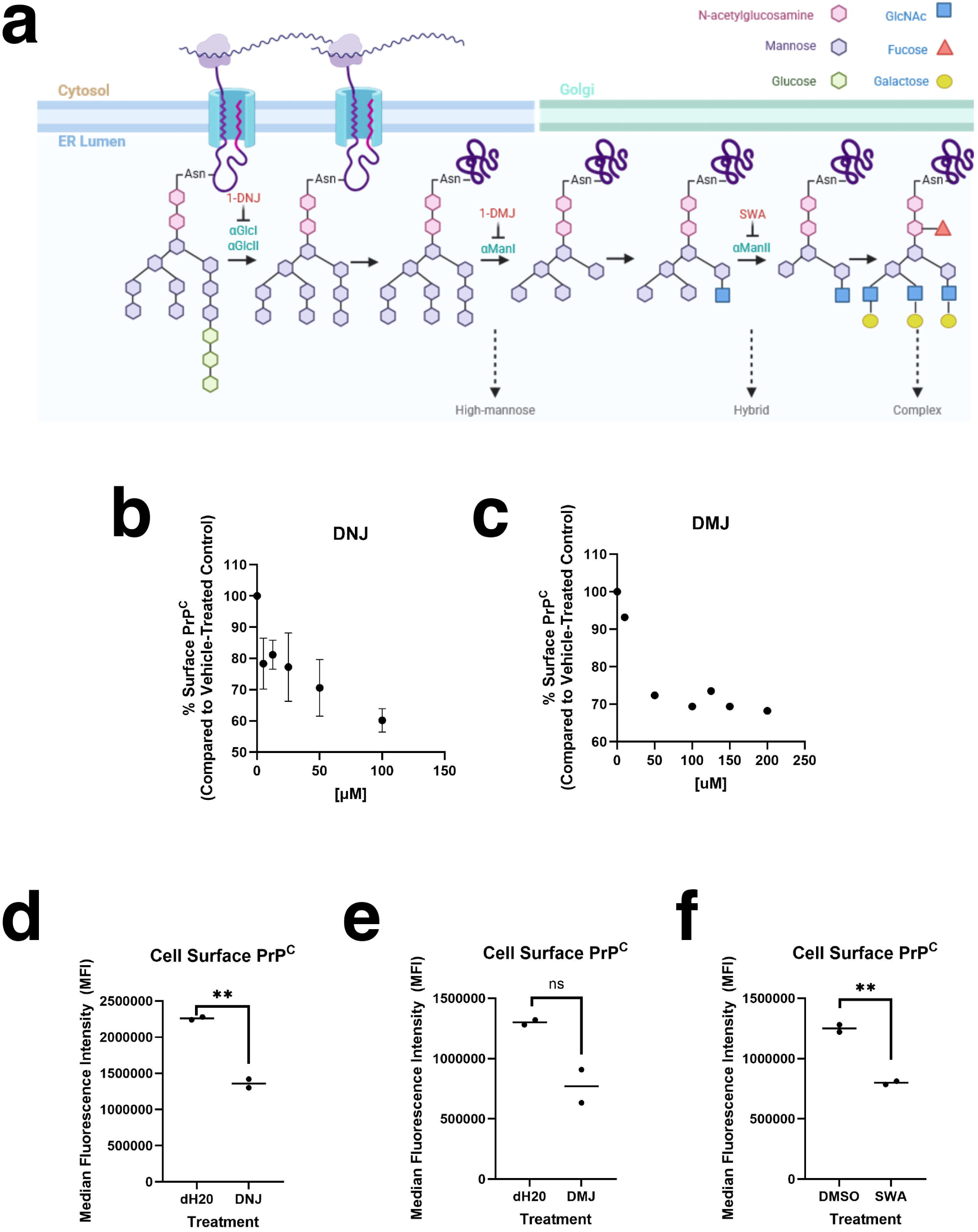
Effects of N-linked glycosylation pathway inhibitors on surface expression of PrP^C^. (A) Schematic showing N-linked glycan biosynthesis in the ER/Golgi. The glycan undergoes a series of glucose and mannose trimming steps before addition of other residues. Enzymes responsible for trimming steps and small molecule inhibitors of these enzymes are labeled. Created in BioRender. Supattapone, S. (2025) https://BioRender.com/42yym9w (B) WT undifferentiated CAD5 cells were treated with varying concentrations of the alpha-glucosidase inhibitor 1-deoxynojirimycin (1-DNJ) in the culture media for 48 hours and cell surface PrP^C^ was measured by flow cytometry. (C) WT undifferentiated CAD5 cells were treated with varying concentrations of the ER specific alpha-mannosidase I inhibitor 1-deoxymannojirimycin (1-DMJ) in the culture media for 24 hours and cell surface PrP^C^ was measured by flow cytometry. (D) Effect of 1-DNJ on cell surface PrP^C^ after 48 hour treatment with 100 µM 1-DNJ or _d_H20 vehicle control as measured by flow cytometry. (E) Effect of 1-DMJ on cell surface PrP^C^ after 72 hour treatment with 50 µM 1-DMJ or _d_H20 vehicle control as measured by flow cytometry. (F) Effect of Golgi-specific alpha-mannosidase II inhibitor swainsonine (SWA) on cell surface PrP^C^ after 72 hour treatment with 10 µM SWA or DMSO vehicle control as measured by flow cytometry. Asterisks represent significance values from unpaired t-tests as follows: *p ≤0.05, **p ≤ 0.01, ***p ≤ 0.001, ****p ≤ 0.0001; n = 2.

Incubation of undifferentiated CAD5 cells with varying doses of the α-glucosidase I/II inhibitor 1-deoxynojirimycin (1-DNJ) **(Figure 4B)** or the ER-specific α-mannosidase I inhibitor 1-deoxymannojirimycin (1-DMJ) **(Figure 4C)** resulted in a dose-dependent decrease in PrP^C^ at the cell surface. Treatment with 1-DNJ **(Figure 4D)**, 1-DMJ **(Figure 4E)**, or the Golgi-specific α-mannosidase II inhibitor swainsonine (SWA) **(Figure 4F)** had similar effects, resulting in approximately 30% reduction in cell surface PrP^C^. None of the treatments had measurable effects on cell morphology **(Supplemental Figure 2A)** or viability **(Supplemental Figure 2B)**. Additionally, the cell surface localization of PrP^C^ was unchanged by drug treatment, as blocking the anti-PrP antibody 6D11 epitope using surface biotinylation **(Supplemental Figure 3A)** or removal of surface PrP^C^ using phosphatidylinositol-specific phospholipase C (PIPLC) **(Supplemental Figure 3B)** resulted in undetectable PrP^C^ levels by Western blot in both drug and vehicle-treated controls. This data confirms our genetic findings and suggests that interfering with the proper N-glycosylation of PrP^C^ and/or other N-glycosylated proteins reduces cell surface levels of PrP^C^ in CAD5 cells.

### Genome-wide knockout screens in differentiated cells reveal shared and unique hits between cell states

CAD5 cells can be reversibly differentiated into a more neuron-like state by the removal of serum from the cell culture medium^79^ **(Supplemental Figure 4A)**. We were interested to see whether the genes responsible for regulating surface PrP^C^ in CAD5 cells would differ in their undifferentiated and differentiated states. Thus, we performed the same genome-wide and secondary library validation screens described above in differentiated CAD5 cells.

In the differentiated state, genome-wide KO screening (in biological triplicate) identified 55 positive regulators of PrP^C^ surface expression and 44 negative regulators of PrP^C^ surface expression (inclusion criteria of log fold change outside 3 standard deviations of the mean (≤ -2.94 or ≥ 2.94) and a *p* value less than or equal to 0.01) **(Figure 5A, Supplementary Tables, “Genome Dif” tab)**. As in the undifferentiated screens, *Prnp* was a top hit (3) **(Supplemental Figure 5A, Supplemental Figure 5B)**. Secondary screening using our custom KO sub-library in differentiated cells validated 41 positive and 13 negative regulators of PrP^C^ cell surface expression (using inclusion criteria of log fold change outside 2 standard deviations of the mean (≤ -1.63 or ≥ 1.63) and *p* value of less than or equal 0.01) **(Figure 5B, Supplementary Tables, “Secondary Val Dif” tab).** Once again, *Prnp* was the top hit.

**Fig. 5.**
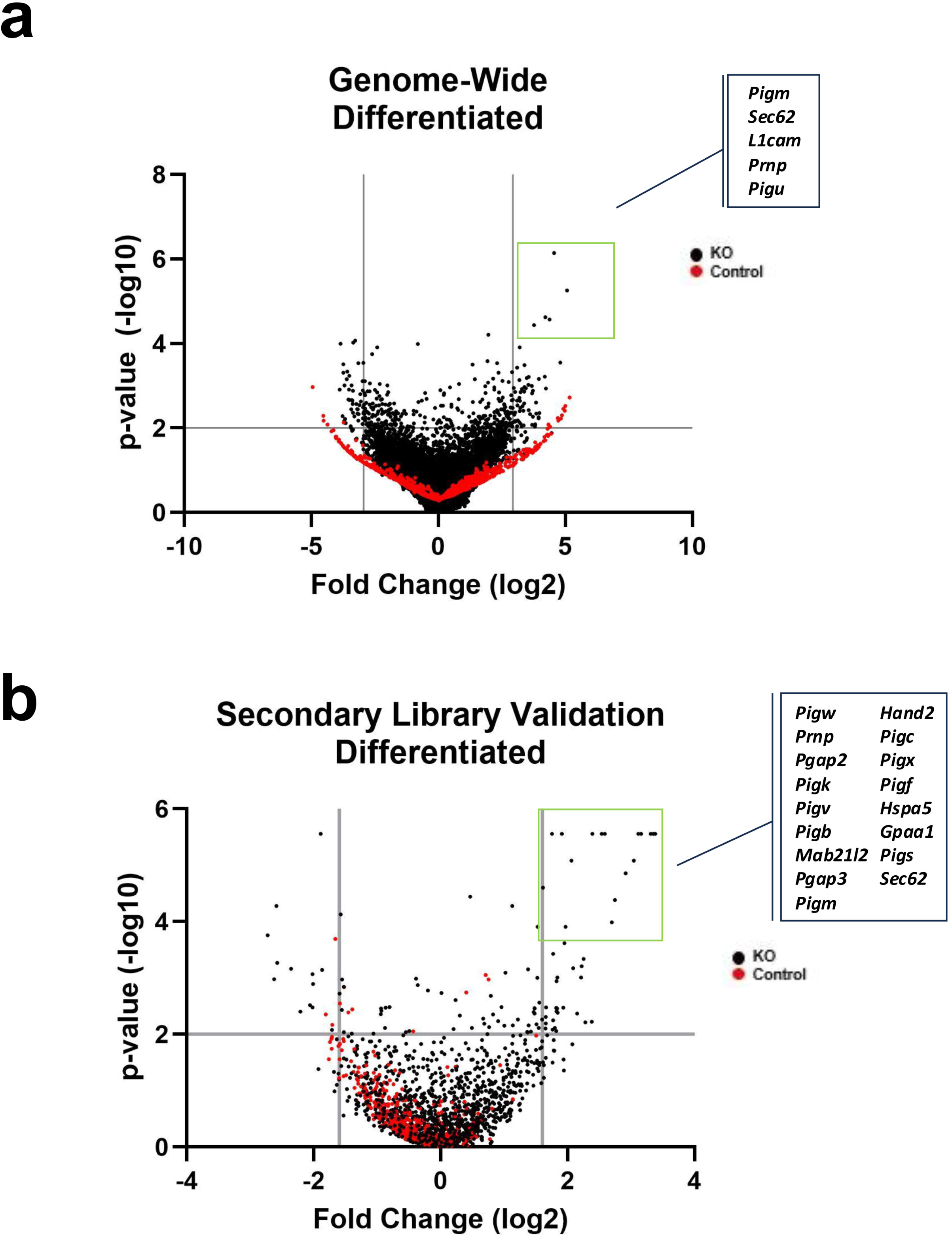
Genome-wide and secondary validation CRISPR-Cas9 screens for positive and negative regulators of PrP^C^ expression at the cell surface in differentiated CAD5 cells. (A) Volcano plot of genome-wide KO screen in differentiated CAD5 cells with top positive regulators of PrP^C^ surface expression labeled. KO genes are plotted as black circles and control genes are plotted as red circles. Cutoff values indicated with gray lines. Cutoffs were -log_10_(*p*-value) ≥ 2 (corresponding to p-value ≤ 0.01) and log fold change ≤ -2.94 or ≥ 2.94 (corresponding to log fold changes outside 3 standard deviations of the mean). (B) Volcano plot of secondary library validation screen in differentiated CAD5 cells with top positive regulators of PrP^C^ surface expression labeled. KO genes are plotted as black circles and control genes are plotted as red circles. Cutoffs were -log_10_(*p*-value) ≥ 2 (corresponding to p-value ≤ 0.01) and log fold change ≤ -1.6 or ≥ 1.6 (corresponding to log fold changes outside 2 standard deviations of the mean).

Top-ranking positive regulators of PrP^C^ surface expression in the undifferentiated and differentiated states were shared, with 8 genes in common between the genome-wide KO screens **(Supplemental Figure 6A)**. Interestingly, no negative regulators of PrP^C^ surface expression were shared between the undifferentiated and differentiated genome-wide screens **(Supplemental Figure 6B)**. Similarly, a greater number of positive regulators of PrP^C^ were shared between the undifferentiated and differentiated cell states during secondary library validation (21 genes) **(Figure 6A)** compared to negative regulators (2 genes) **(Figure 6B)**.

**Fig. 6.**
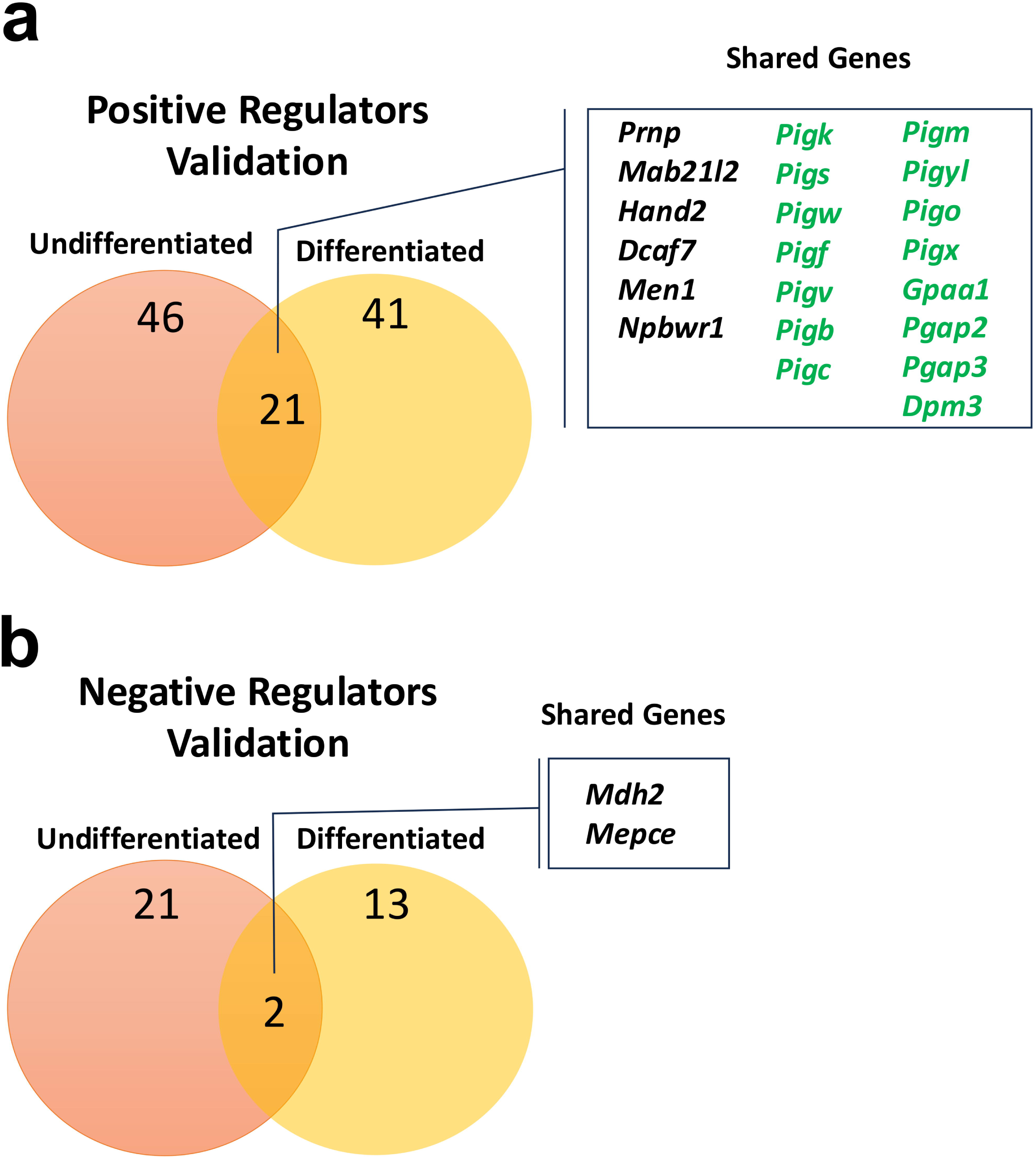
Comparison of genetic and pathway hits between undifferentiated and differentiated CAD5s. (A) Visual diagram comparing the number of positive regulator gene hits identified in the undifferentiated and differentiated secondary validation screens, with shared genes labeled. Genes which belong to the GPI-anchor biosynthesis pathway are labeled in green. Other genes are labeled in black. (B) Visual comparison of the number of negative regulators of cell surface PrP^C^ identified in the validation screens in the undifferentiated and differentiated states. Shared genes are labeled.

Secondary validation screening revealed that most genetic regulators of cell surface PrP^C^ were dependent on cell state **(Figure 6A**, **Figure 6B)**, including the N-linked glycosylation pathway genes which were unique to the undifferentiated state **(Figure 3B, purple)**. However, our studies also identified core regulatory genes and pathways which were shared between the undifferentiated and differentiated cell states, including many GPI-anchor biosynthesis pathway genes **(Figure 6A -green text)**. Taken together, our studies identify unique genes which regulate cell surface PrP^C^ in a state-dependent manner, as well as core shared genes and pathways which represent fundamental regulators of cell surface PrP^C^ expression in a model system with direct relevance to disease.

### Hspa5 inhibitor reduces PrPC surface expression in undifferentiated and differentiated CAD5 cells

Several genes encoding heat shock protein 70 (HSP70) family members were identified as hits in our screens. HSP70 family members are involved in the folding of newly synthesized proteins, refolding of misfolded and aggregated proteins, membrane translocation of secretory proteins (reviewed in^89^) and have previously been studied in relation to prion disease^90–93^. Therefore, we were interested in testing the effect of a small molecule inhibitor of Hspa5 (also known as BiP) on cell surface PrP^C^ levels in undifferentiated and differentiated CAD5 cells. Hspa5 inhibition using the inhibitor HM03 caused a decrease in cell surface PrP^C^ expression in both undifferentiated **(Supplemental Figure 7A)** and differentiated **(Supplemental Figure 7B)** CAD5 cells with no apparent effects on the cellular localization of the remaining PrP^C^ **(Supplemental Figure 7C)**. This data confirms our genetic findings and suggests that interfering with the function of Hspa5 reduces cell surface levels of PrP^C^ in CAD5 cells.

## Discussion

We present the first genome-wide CRISPR/Cas9 KO screen for regulators of PrP^C^ cell surface expression in a prion-infectible cell line of neuronal origin, CAD5. Intriguingly, our results revealed both similarities and differences between undifferentiated and differentiated CAD5 cells, suggesting that some regulatory pathways are dependent on cell state.

### CAD5 cells display similarities and differences in PrP^C^ regulation mechanisms relative to non-infectible cell types

Two genome-wide screens for modulators of PrP^C^ have been previously published. Davis et al., utilized gene-trap mutagenesis in human haploid myeloid HAP1 cells to compare genes which regulate surface expression of two different GPI-anchored proteins, PrP^C^ and CD59^60^. Heinzer et al., utilized RNAi to achieve gene knockdowns in human U251-MG glioblastoma cells and identified genes which altered total PrP^C^ levels in whole cell lysate^36^. We were interested in identifying genes which regulate PrP^C^ levels at the cell surface and thus, our genome-wide screens utilized CRISPR/Cas9 to achieve complete gene knockouts in prion-infectible cells and assayed for PrP^C^ abundance at the surface of live CAD5 cells.

Our screens identified numerous hits related to PrP^C^ biosynthesis and trafficking (including GPI-anchor biosynthetic and N-glycan biosynthetic genes). This is in stark contrast to the RNAi screens by Heinzer et al., which did not identify GPI-anchor or N-glycan biosynthetic hit genes^36^. This could be due to differences in cell type but is more likely attributable differences in screening phenotype (whole cell lysate PrP^C^ vs. cell surface PrP^C^), highlighting the strength of our approach. We validated 15 GPI-anchor biosynthetic genes as shared positive regulators of PrP^C^ cell surface expression in undifferentiated and differentiated CAD5 cells. 10 of these genes were previously identified by Davis et al. as important for regulation of cell surface PrP^C^ in HAP1 cells^60^, suggesting that the GPI-anchor biosynthetic pathway is important for PrP^C^ cell surface expression in multiple cell types, importantly, including prion-infectible CAD5 cells. Two of these 15 GPI-anchor biosynthetic genes, *Pigf* and *Pgap2* were validated as positive regulators of cell surface PrP^C^ expression in our undifferentiated and differentiated CAD5s screens but were identified as genes unique to the regulation of CD59 cell surface expression (not regulators of PrP^C^ cell surface expression) by Davis et al. in HAP1 cells. Additionally, we identified *Pigyl* (the mouse equivalent of human PIGY) as a hit, while Davis et al. did not. It is unknown whether these discrepancies are due to differences in cell type, screening methodology, or statistical inclusion limits for hit calling. It is unclear modulation of GPI anchoring would be a viable therapeutic strategy as prior studies show that anchorless PrP^C^ is secreted and can form infectious prions, but is inefficient at transducing toxicity to the host^94–96^.

The PrP precursor polypeptide undergoes cotranslational translocation and enters the ER via the Sec61 protein-conducting channel but is dependent on Sec62/Sec63 for stabilization due to its lack of structured domains in the N-terminal region^97–99^. Davis et al. identified *Sec62* and *Sec63* as positive regulators of PrP^C^ cell surface expression in HAP1 cells^60^. In our genome-wide screens, *Sec62* was identified as positive regulator of CAD5 PrP^C^ surface expression in both the undifferentiated state and differentiated states. However, *Sec62* only met the hit inclusion criteria for the secondary validation screens in the differentiated cell state. Surprisingly, *Sec63* was not identified as a regulator of CAD5 PrP^C^ cell surface expression in any of our screens. This could potentially be explained by the fact that Sec63’s role is substrate-specific^97^ and that its function can partially be compensated for by other proteins, particularly by other J-domain proteins which can activate the ATP-ase function of Hspa5 (also known as binding immunoglobulin protein, BiP)^100^.

Interestingly, *Hspa5* was validated as a positive regulator of PrP^C^ cell surface expression in the differentiated cell state but was not identified in either of the previously published genome-wide screens for PrP^C^ regulators^36,60^. *Hspa5* encodes for the Hspa5 protein which serves as a molecular chaperone in many contexts. Hspa5 facilitates Sec61 gating for PrP entry to the ER lumen for cotranslational translocation^97–99,101^. It also promotes the correct folding of PrP^C^^90,91^. Thus, our identification of *Hspa5* as a positive regulator of cell surface PrP^C^, further validated using a small molecule inhibitor, highlights its multiple roles in PrP^C^ biosynthesis and regulation. Interestingly, Hspa5 also plays a role in prion pathogenesis, as reduced expression of Hspa5 has been shown to accelerate prion pathogenesis *in vitro* and *in vivo*^102^. This acceleration of prion pathogenesis may relate to its key role in maintaining protein homeostasis via activation of the unfolded protein response (UPR)^91^, illustrating the complexities of secretory protein biosynthesis, folding, and protein homeostasis in relation to prion disease.

Our genome-wide screens did not identify several PrP^C^ regulators previously identified by Heinzer et al. including *Nxf1, Kansl1, Tmcc2, Aplp2, Pum1, Puf60*^36^. We purposefully included these genes in our secondary validation library to give them an additional opportunity to be identified as hits, but none reached our threshold criteria for inclusion as a hit. This may be due, in part, to the differences in genetic perturbation method used (RNAi knockdown vs. CRISPR/Cas9 knockout), cell type and species (human U-251 MG vs. mouse CAD5), or screening phenotype and methods (whole cell lysate ELISA vs. live cell surface FACS).

Despite these differences, we identified a handful of identical hits to Heinzer et al. in our genome-wide and/or secondary library screens including *Ap5s1, Copg1, and Copz1*^36^. Additionally, we identified hit genes encoding proteins from the same multi-protein complexes identified by Heinzer et al., including many spliceosome-related genes (*Srsf3, Snrnp200, Snrpd3, Sf3b3, Sf3b2* and *Sf3b1*)^103^ and the transcription export (TREX) complex member *Thoc6*^104^. Interestingly, Heinzer et al. note that 4 of their 9 validated PrP^C^ regulators (*Copg1, Copz1, Sf3a1, and Puf60*) also seem to affect glycosylation of PrP^C^, which may further support our finding that N-glycosylation regulates cell surface PrP^C^ expression.

### Some mechanisms regulating cell surface PrP^C^ in CAD5 cells are dependent on cell state

Key differences in the cell surface PrP^C^ regulatory pathways of undifferentiated and differentiated CAD5 cells were identified by our genetic screens and may contribute to our understanding of the complex interplay between PrP^C^ expression, cell state determination/differentiation, and prion infectibility.

In undifferentiated CAD5 cells, several genes related to the N-glycosylation biosynthetic pathway (including *Mgat1*, *Alg5*, *Alg10b*) were validated as positive regulators of cell surface PrP^C^. Small molecule inhibitors of N-glycosylation pathway enzymes were used as an additional method of validating our finding that N-glycosylation influences cell surface expression of PrP^C^. However, it is worth noting that no inhibitors of the specific hit N-glycosylation genes identified in our study were tested, rather we focused on validating the pathway in general. That the hit N-glycosylation genes were identified only in the undifferentiated state may reflect differences in the N-glycome between undifferentiated and differentiated cells as well as differences in protein quality control and homeostasis mechanisms between cell states^105–112^. Interestingly, the N-glycosylation of PrP^C^ has been studied extensively in relation to its effects on PrP^C^ localization^113–116^, role in limiting PrP^Sc^ transmission between species^117–119^, and PrP^Sc^ infectivity^115,118–134^.

In differentiated CAD5 cells, unique positive regulators of the cell surface expression of PrP^C^ largely related to cellular processes downstream of transcription, including genes related to protein trafficking (*Ap1m2, Tmed10*) and protein folding in response to cellular stress (*Hspa5*, *Gsta2*)^135–139^. Interestingly, several genes related to synaptic function and neurodegenerative disease were also validated as positive regulators of cell surface PrP^C^ expression solely in the differentiated state (including *Siva1*, *Unc13a,* and *Gsk3b*). *Siva1* encodes an adaptor protein that regulates synaptic function through its interaction with the E3 ubiquitin ligase XIAP and plays a role in receptor internalization in synapses^140^. *Unc13a* encodes a key regulator of synaptic vesicle priming and neurotransmitter release and variants in *Unc13a* are linked to ALS and frontotemporal dementia (FTD) (reviewed in^141^). *Gsk3b* encodes a serine/threonine kinase implicated in Alzheimer’s disease (reviewed in^142^).

Negative regulators of the cell surface expression of PrP^C^ also largely differed in identity between cell states but were functionally much more closely related. Negative regulators in both cell states were primarily related to gene expression regulation including RNA processing and transcription factors. Negative regulators validated in the undifferentiated cell state also related to DNA maintenance (*Chek1*, *Ankrd17*)^143,144^, while those in the differentiated state did not, potentially reflecting differences in cell cycle between actively proliferating and quiescent cells.

Taken together, the fact that certain genes were identified solely in undifferentiated or differentiated cells suggests that some mechanisms responsible for regulating cell surface PrP^C^ are sensitive to cell state and underscores the importance of performing genetic discovery experiments directly in neuronal-like prion-infectable cells relevant to disease.

Technical limitations could also potentially explain the differences in gene hits between undifferentiated and differentiated CAD5 cells. We wondered if some genes were only identified in one cell state because knockout of certain genes could impact the differentiation process or viability. Use of a viability dye in our FACS process could have served as a control for potential differences in viability but was omitted. We compared the abundance of sgRNAs targeting such genes in the undifferentiated and differentiated unsorted library control populations. We observed that the sgRNAs targeting hit genes from one or the other cell state were still present at relatively equal proportions in each unsorted library control, indicating that loss of cells containing specific knockouts due to viability issues was not likely to be causing the differences in hit genes between cell states. However, we observed that even CAD5 cells considered to be fully differentiated displayed a high degree of morphological diversity, as previously described^145^. It is likely that our differentiated screens encapsulated cells at different points on the spectrum of differentiation. Similar amounts of cell surface PrP exist in fully differentiated CAD5s and fully undifferentiated (proliferative) cells^75^, however, cells with different differentiation statuses are likely to have different gene expression profiles and transcriptional activity. The heterogeneity in differentiation state, differences in transcriptional activity and differences in gene expression may have contributed to the lower significance values and higher false discovery rates observed in our differentiated screens compared to the undifferentiated screens and may have impacted our identification of hit genes.

### Some core pathways and genes regulate PrP^C^ expression in CAD5 cells independent of differentiation state

We validated 21 positive regulators and 2 negative regulators of surface PrP^C^ as shared hits between the undifferentiated and differentiated cell states. Most of the shared positive regulators (15 genes of 21 genes) fell within the GPI-anchor biosynthetic pathway. These hits were unsurprising, as PrP^C^ is a GPI-anchored protein and thus relies on the GPI-anchor fo proper tethering at the cell surface^11,146,147^. The remaining shared positive regulators of PrP^C^ cell surface expression in undifferentiated and differentiated CAD5 cells span the lifecycle of PrP^C^, including genes encoding PrP itself (*Prnp*), a chromatin associated protein (*Men1*)^148^, transcription factors (*Mab21l2* and *Hand2*)^149–151^, protein quality control and degradation (*Dcaf7*)^42,152,153^ , and a neuropeptide receptor (*Npbwr1*)^154^. The shared negative regulators of PrP^C^ surface expression in the undifferentiated and differentiated cell states encode a metalloprotease (*Mepce*)^155^ and an enzyme involved in the citric acid cycle (*Mdh2*)^156^.

One interesting observation about the shared hits between the undifferentiated and the differentiated state was that the discrepancy in number of positive regulatory gene hits (21 hits) in comparison to the number in the negative regulatory hits (2 hits). An explanation for this difference could have to do with the degree of variability in PrP^C^ expression after gene perturbation. Our FACS gating scheme purposefully included cells with no or low PrP^C^ abundance at the cell surface (positive regulators of PrP^C^). A gene knockout that results in a binary signal (no surface PrP^C^) would be expected to result in a complete absence of fluorescent signal in each cell containing the knockout. Thus, all cells with the gene knockout would be expected to be captured in the low PrP^C^ population, resulting in a high significance value. However, genetic perturbations that cause an increase in surface PrP^C^ would likely be more variable, with less consistency between cells containing the same gene knockout and lower significance values as a result. An additional contributor to the unevenness in hit genes between the positive and negative regulators may relate to essentiality or lethality. Perhaps most genes that negatively regulate PrP^C^ also negatively regulate other proteins, which, when unregulated, lead to cell stress or death.

While we believe the genetic regulators of PrP^C^ surface expression in CAD5 cells identified through our genome-wide screening efforts are useful, we do acknowledge the limitations of the study. We identified genes which reduce cell-surface expression of PrP^C^ in a prion-infectible cell line, but we did not directly test whether modulation of the identified genes influences prion infectibility or propagation. Additionally, CRISPR/Cas9 pooled genetic screens may not fully capture functionally redundant genes, genes with overlapping function, essential genes, or genes which impact cell viability and some genes which were incompletely knocked out or genes with low abundance guides may have been lost due to selection bias. We utilized *p* values and log fold change values from MAGeCK analysis to identify inclusion criteria for hit genes. Using different bioinformatic analysis packages or setting different inclusion criteria may alter the ranking and number of hit genes. For this reason, we have deposited all raw .fastq files from our primary and secondary library screens for further analysis by interested parties on Mendeley Data.

## Conclusions

Our unbiased genome-wide CRISPR/Cas9 KO screens provide a canonical list of genes which regulate PrP^C^ expression on the surface of neuronal-like undifferentiated and differentiated CAD5 cells. We identified and validated 21 core positive regulators and 2 negative regulators which were shared between cell states. Surprisingly, some genes were only identified in the undifferentiated or differentiated cell states, suggesting that some mechanisms for PrP^C^ regulation may be state-dependent, highlighting the importance of identifying regulators in cell models relevant to disease. This core list of genes responsible for regulating cell surface PrP^C^ expression in a highly prion-susceptible, neuron-like cell type provides a useful roadmap for future studies and may provide insight into potential therapeutic targets for prion disease and other neurodegenerative diseases.

## Notes

doi:10.17632/jv32cvpn3r.1

