## Supplementary Figures for "Genome-Wide Screens Identify Core Regulators of Cell Surface Prion Protein Expression"

**Genome-wide knockout screens identify regulators of cell surface prion protein expression in prion-susceptible CAD5 cells**

Kathryn S. Beauchemin^1^ and Surachai Supattapone^1,2,*^

Departments of Biochemistry and Cell Biology^1^ and Medicine^2^, Geisel School of Medicine at Dartmouth, Hanover, New Hampshire 03755, USA

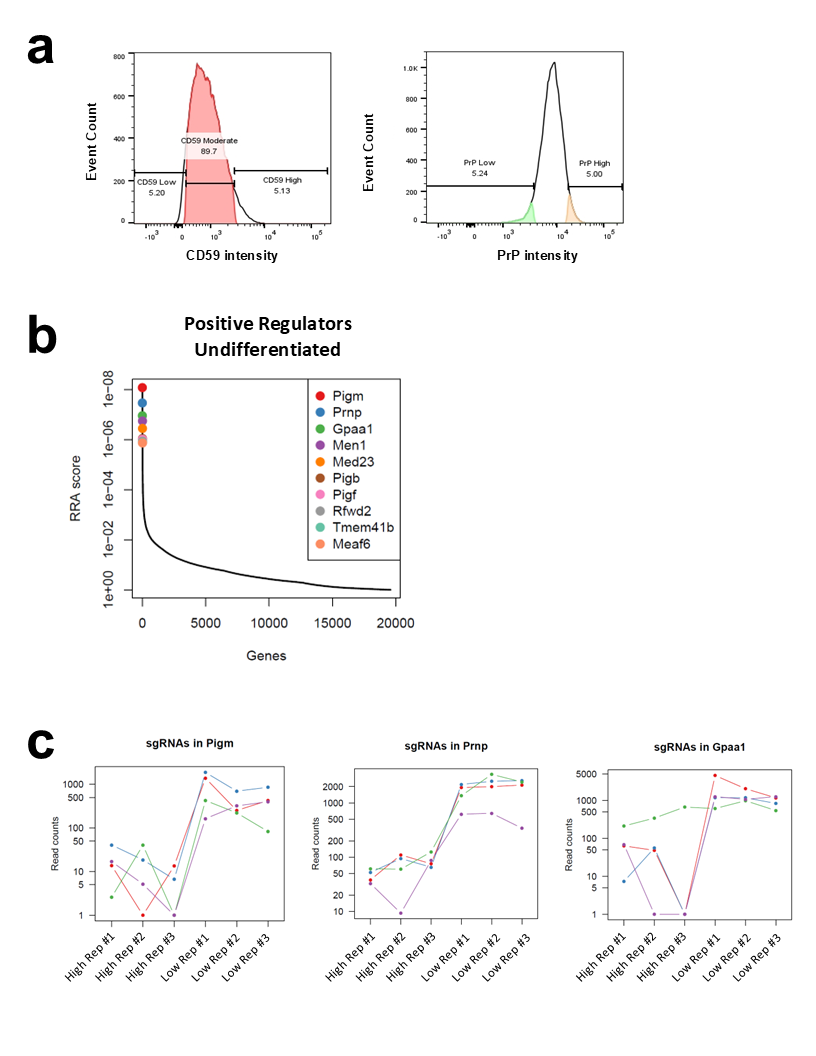

**Supplemental Fig. 1.** (A) Representative flow cytometry histograms of CAD5 genome-wide library immunostained with anti-CD59:APC (left panel) and anti-PrP primary plus anti-mouse secondary:PE (right panel) for FACS. (B) MAGeCK-VISPR analysis of NGS results from the whole-genome KO screen for PrP^C^ cell surface expression in undifferentiated CAD5 cells were plotted as a distribution of RRA values across genes, with the top ten positive regulator hits highlighted. (C) Read count plots from three of the top ranked hits in the genome-wide KO screens in undifferentiated CAD5 cells showing agreement in enrichment of guides within “PrP^C^ Low” samples as compared to “PrP^C^ High” samples. Each line represents one sgRNA.

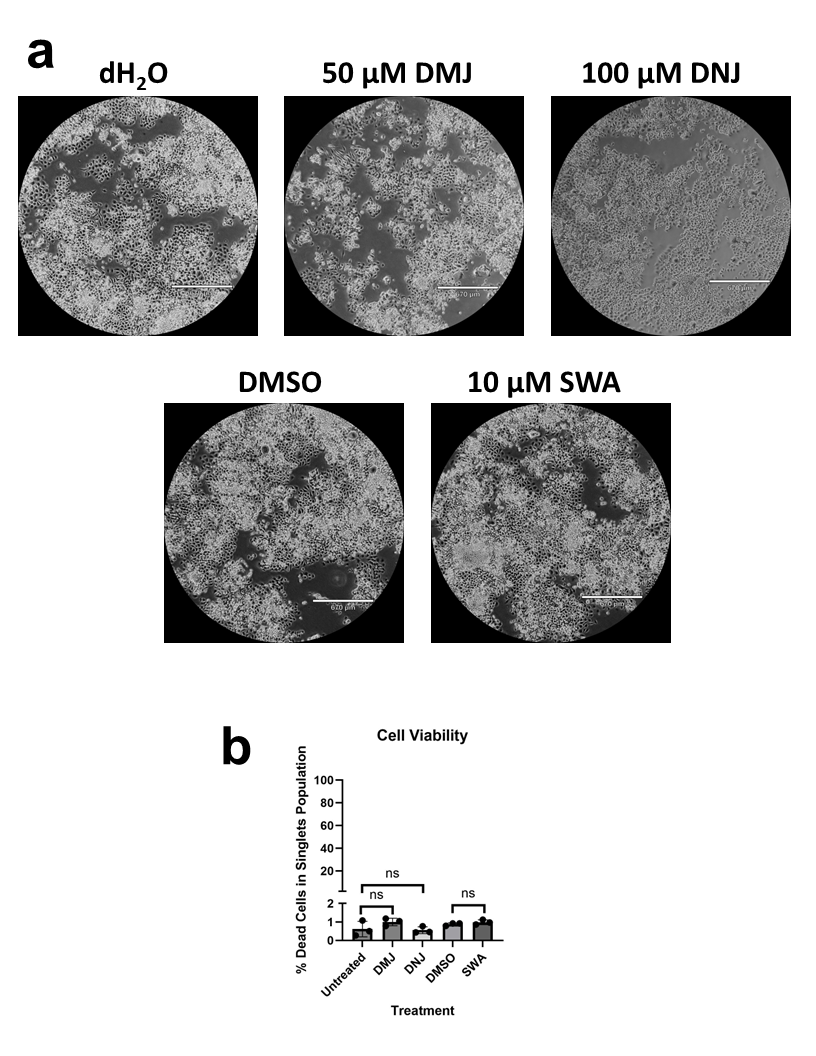

**Supplemental Fig. 2.** (A) Representative microscopy images of WT CAD5 cells treated N-glycosylation inhibitors for 72 hr at indicated concentrations under 40X magnification. (B) Plot showing percentage of dead cells in single cell populations as labeled by membrane impermeant viability dye via flow cytometry after 72 hr treatment with N-glycosylation inhibitors. Asterisks represent significance values from unpaired t-tests as follows: ns = no significance.

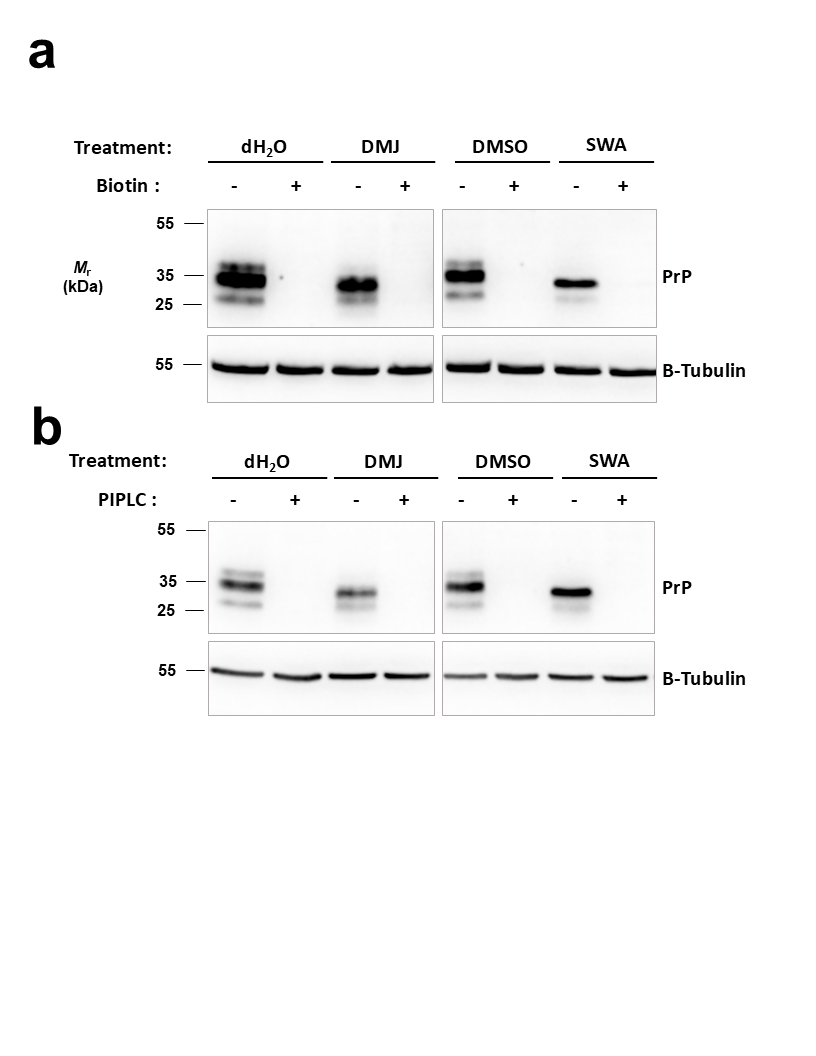
**Supplemental Fig. 3.** Western blot showing reduction in total PrP^C^ in WT CAD5 whole cell lysate after 72 hr treatment with N-glycosylation inhibitors. Drug-treated or vehicle-treated PrP^C^ remains localized to the cell surface, as (A) abrogation of anti-PrP antibody epitope for binding to cell surface PrP^C^ via surface biotinylation or (B) enzymatic release of phosphatidylinositol-linked proteins using phosphatidylinositol-specific phospholipase C (PIPLC) results in complete loss of detectable PrP^C^.

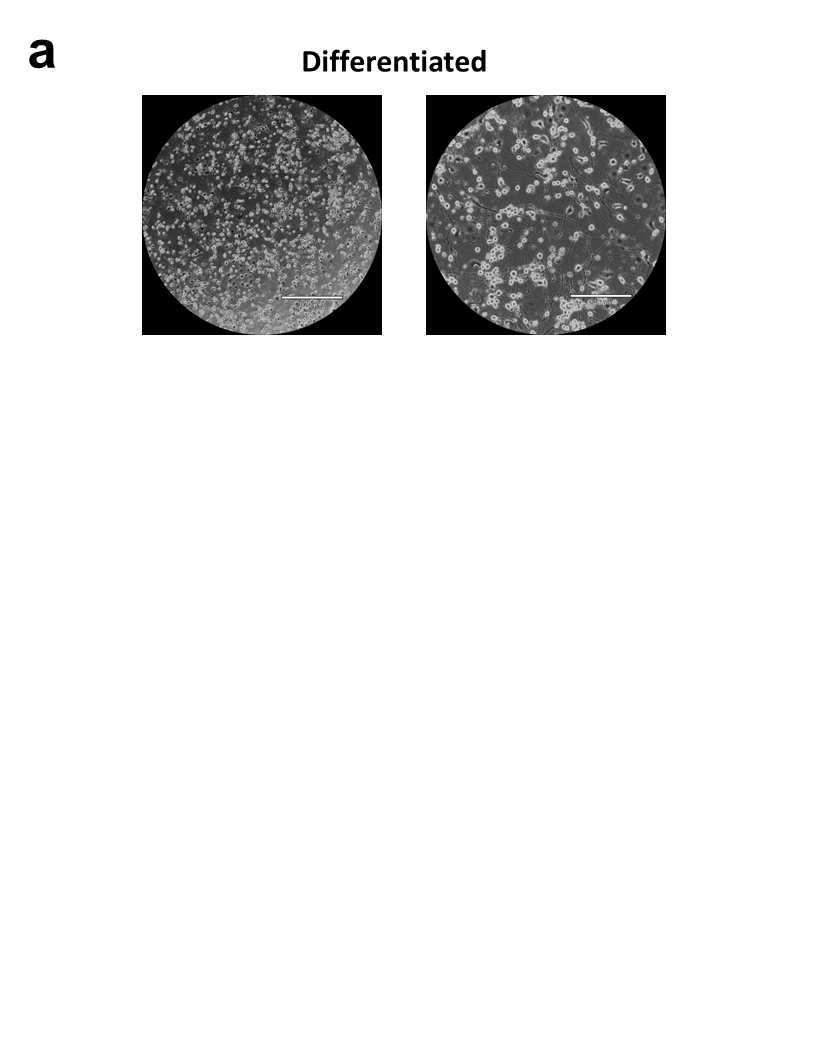

**Supplemental Fig. 4**. (A) Representative microscopy images showing growth of neuronal processes in differentiated WT CAD5 cells after serum deprivation in protein-free media for 96 hr at 40X (left) or 100X (right) magnification.

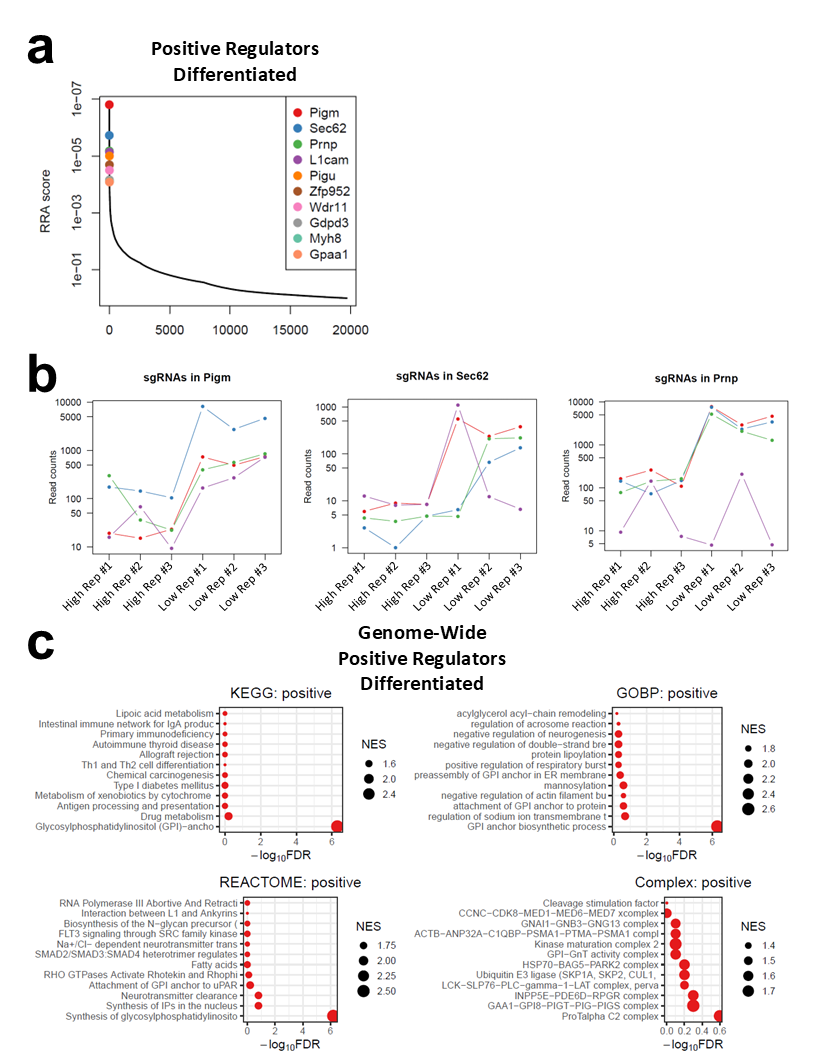

**Supplemental Fig. 5.** (A) MAGeCK-VISPR analysis of NGS results from the whole-genome KO screen for PrP^C^ cell surface expression in differentiated CAD5 cells were plotted as a distribution of RRA values across genes, with the top ten positive regulator hits highlighted. (B) Read count plots from three of the top ranked hits in the genome-wide KO screens in differentiated CAD5 cells showing agreement in enrichment of guides within “PrP^C^ Low” samples as compared to “PrP^C^ High” samples. Each line represents one sgRNA. (C) Pathway analysis of genome-wide differentiated positive hits was conducted using the Kyoto Encyclopedia of Genes and Genomes (KEGG)^1-3^, Gene Ontology Biological Process (GOBP), Reactome, and Complex databases through the MAGeCKFlute platform. Negative log_10_ false discovery rates (FDR) were plotted for pathway hits and point size corresponds to relative normalized enrichment score (NES).

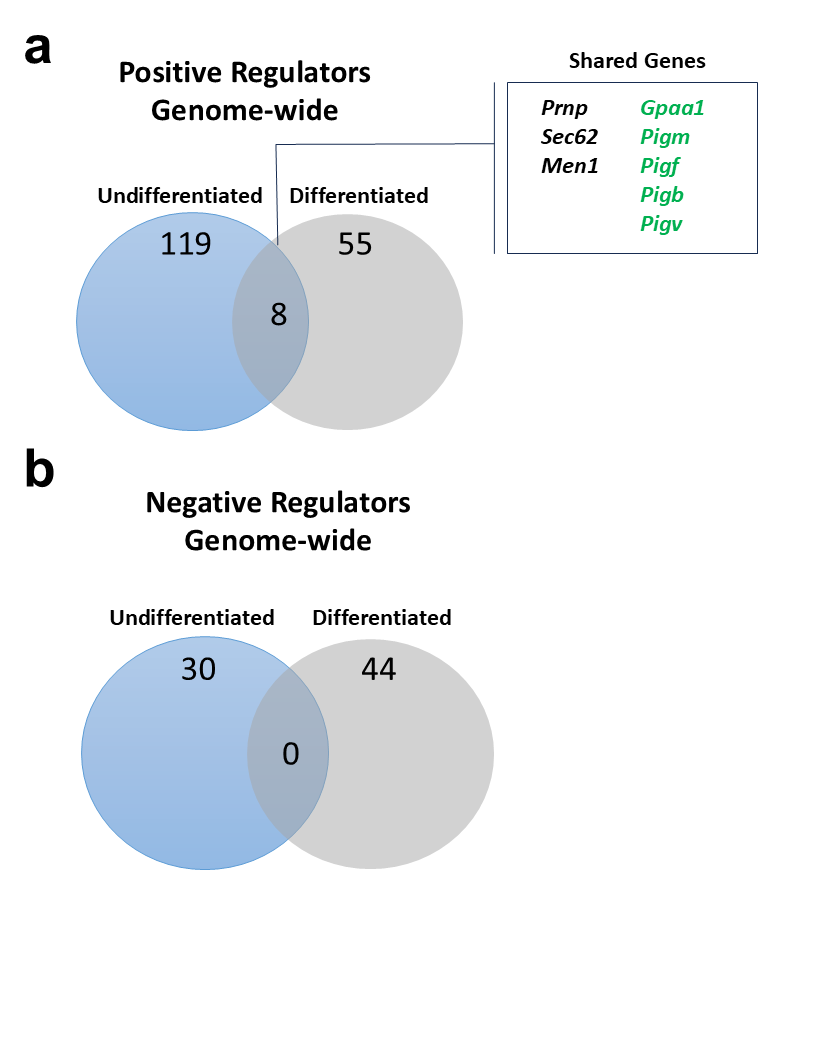
**Supplemental Fig. 6.** (A) Visual comparison of the number of positive regulators of PrP^C^ surface expression identified in the genome-wide screens in the undifferentiated and differentiated states. Shared genes are labeled, with those relating to GPI-anchor biosynthesis labeled in green. (B) Visual comparison of the number of negative regulators of PrP^C^ surface expression identified in the genome-wide screens in the undifferentiated and differentiated states.

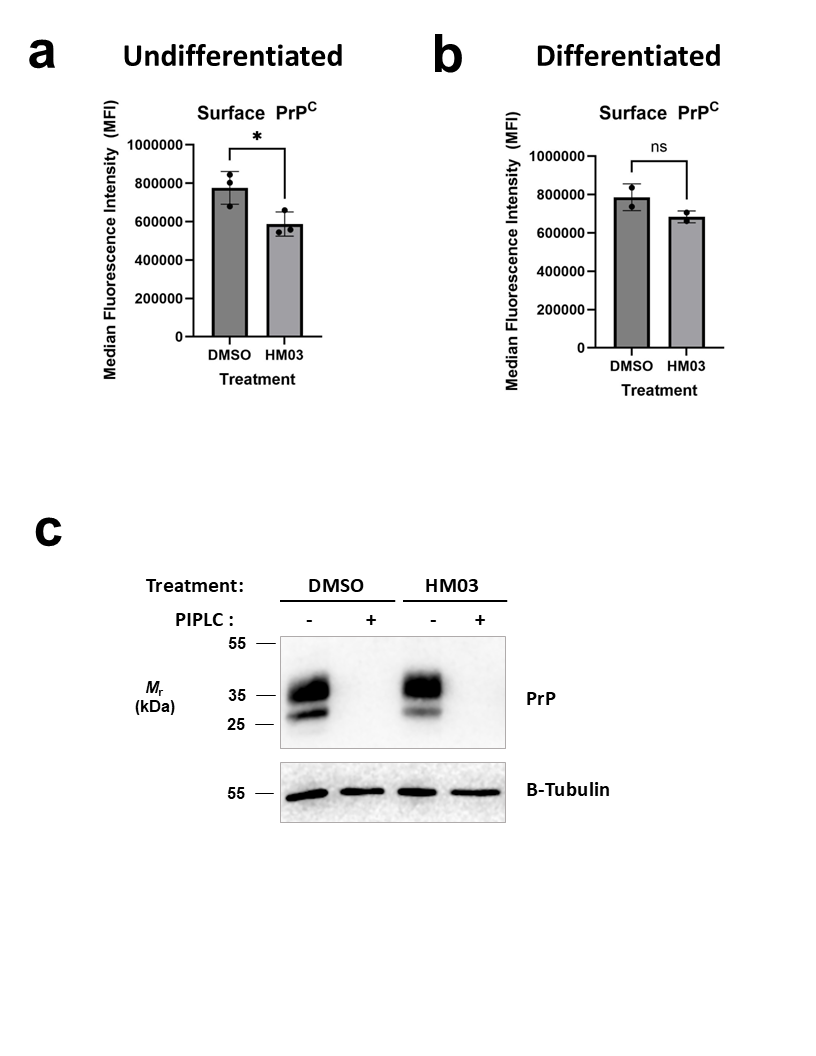
**Supplemental Fig. 7.** WT CAD5 cells were treated with 5 µM HM03 (Hspa5 inhibitor) in the culture media for 24 hr and cell surface PrP^C^ was measured by flow cytometry. (A) Effect of HM03 on cell surface PrP^C^ in undifferentiated CAD5 cells after 24 hr treatment with 5 µM HM03 or DMSO vehicle control as measured by flow cytometry. (B) Effect of HM03 on cell surface PrP^C^ in differentiated CAD5 cells after 24 hr treatment with 5 µM HM03 or DMSO vehicle control as measured by flow cytometry. Asterisks represent significance values from unpaired t-tests as follows: *p ≤0.05, ns = no significance. (C) Western blot showing reduction in total PrP^C^ in undifferentiated WT CAD5 whole cell lysate after 24 hr treatment with the Hspa5 inhibitor, HM03. Drug-treated or vehicle-treated PrP^C^ remains localized to the cell surface, as enzymatic release of phosphatidylinositol-linked proteins using phosphatidylinositol-specific phospholipase
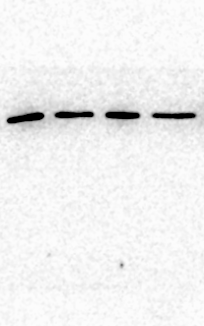

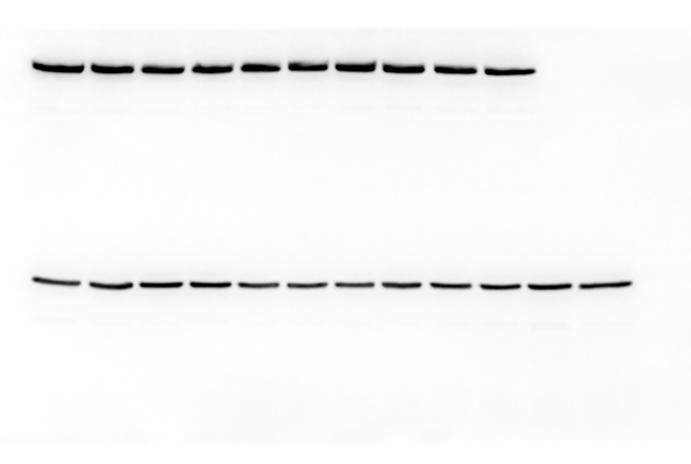

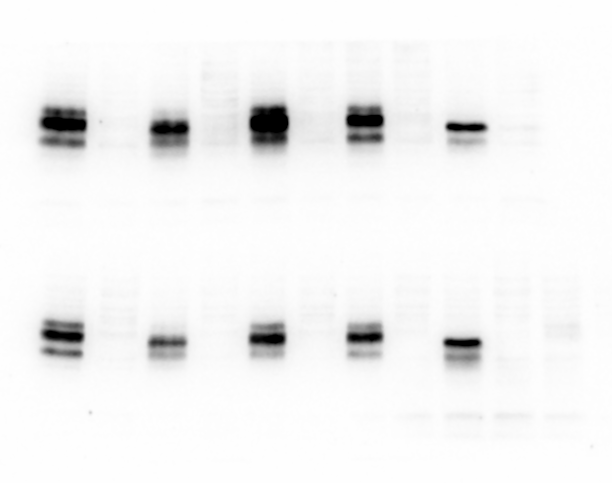
C (PIPLC) results in loss of detectable PrP^C^.

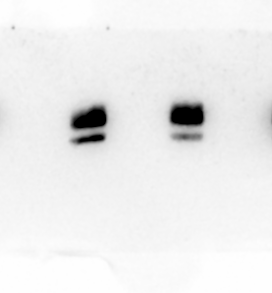

**Supplemental Fig. 8.** Uncropped images for Western blots show in Supplemental Figures 3 and 7C.
